# Hypergraph geometry localizes mutation-associated dependency vulnerabilities in cancer

**DOI:** 10.64898/2026.07.26.740812

**Authors:** Jiyun Lee, Jihyo Lee

## Abstract

Mutation-specific therapeutic vulnerabilities remain difficult to identify in precision oncology because lineage effects and network topology can obscure true synthetic lethal relationships. Here, we present a computational framework that maps genomic mutational profiles and genome-wide CRISPR-Cas9 screens onto multi-scale hypergraphs constructed from macromolecular complexes (CORUM), protein interaction modules (STRING), and transcriptional regulons (TRRUST). Using strict reproducibility criteria across lineage-split cohorts, we identify cross-topology overlap families of dependencies that recur across independent biological organizational layers. We then characterize these candidates with discrete hypergraph geometry, including Hypergraph Fractional Ricci Curvature (HFRC) and Hypergraph Local Ricci Curvature (HLRC), and evaluate them against multidimensional matched-null distributions and degree-preserving shuffles to control for network density and node degree bias. We evaluate the 8 core cross-topology families for clinical prognostic value. Separately, to validate the pharmacological actionability of the hypergraph framework, we project our prioritized candidates onto PRISM dose-response profiles, confirming that the geometrically constrained BRAF:MAP2K1 family exhibits strong, selective drug sensitivity to downstream MAPK pathway inhibition. The strongest face-validity result is SMARCA2:SMARCA4, a clinically validated paralog synthetic lethal pair in chromatin remodeling that our pipeline recovered purely through unsupervised geometric constraints. While across the prioritized overlap families this specific class of reproducible dependencies localizes to highly integrated, negatively curved hypergraph regions, this topological separation is statistically fragile; it achieves nominal significance (*p* = 0.031) as a primary endpoint and is sensitive to the removal of single families. Nonetheless, this pattern persists across independent screening platforms, including Sanger and DepMap 25Q3, suggesting a potential, albeit statistically sensitive, geometric framework for prioritizing actionable targets in precision oncology.

## 1. INTRODUCTION

The primary objective of precision oncology is to identify therapeutic vulnerabilities that are selectively lethal to cancer cells harboring specific genomic alterations while sparing normal tissue [46]. Decades of cancer sequencing have generated exhaustive catalogs of somatic alterations, categorized broadly into damaging mutations (loss-of-function variants like truncations or frameshifts) and hotspot mutations (putative gain-of-function or dominant-negative alterations) [47]. Throughout this study, analyzed mutation classes include both recurrent hotspot alterations and predicted damaging variants. However, translating these complex genomic signatures into clinical targets remains a formidable challenge [3]. While a subset of oncogenic drivers can be directly targeted with small-molecule inhibitors, many mutations reside in proteins that have historically been considered undruggable [4]. Consequently, research has shifted toward uncovering mutation-specific genetic dependencies, such as synthetic lethality or collateral vulnerabilities, wherein a somatic alteration induces selective dependency on a partner gene [5, 6].

Large-scale, high-throughput functional genomics platforms have revolutionized target discovery [7]. Computational and experimental methods increasingly combine clinical networks with CRISPR perturbations to systematically prioritize synergistic drug targets and functional pathways [8]. Initiatives such as the Cancer Dependency Map (DepMap) utilize genome-wide CRISPR-Cas9 screens across hundreds of molecularly characterized cancer cell lines to map gene essentiality [9]. High-resolution genomics arrays, such as Project DRIVE [10], provide complementary multidimensional loci of cancer cell dependencies, while multi-omic profiles from the Cancer Cell Line Encyclopedia (CCLE) provide key baseline genetic annotations for evaluating cohort stratification [11]. However, translating screen-derived associations into robust therapeutic hypotheses remains challenging because many candidate dependencies exhibit context-specificity across lineages, datasets, or perturbation platforms [12, 13]. This context-specificity is frequently driven by statistical false positives, lineage-associated confounding (where passenger alterations and unrelated dependencies are co-enriched in specific tissue origins), and heterogeneous network connectivity [13]. To mitigate these confounding factors, reproducibility across independent biological contexts must be treated as a primary design constraint in computational discovery pipelines [12]. Accordingly, candidate dependencies were required to demonstrate reproducibility across independently stratified biological contexts rather than being inferred from a single pooled analysis. Furthermore, existing network-constrained approaches typically rely on a single biological network representation, making it difficult to distinguish robust dependencies from topology-specific artifacts [14, 15].

To address these limitations, network-constrained frameworks leverage known molecular interactions to filter out spurious, non-physical correlations [16]. In particular, network-based inference methods have proven successful in mapping the functional impact of somatic alterations onto transcriptional regulators and signaling cascades [17]. However, traditional approaches model cellular architectures using binary pairwise graphs [14]. In reality, biological systems are fundamentally multi-component; proteins function collectively within higher-order macromolecular complexes and transcriptional regulatory units [18]. Rather than relying on flat graph abstractions, higher-order physical and regulatory assemblies can be represented as hypergraphs, capturing complex non-pairwise dynamics that are lost in traditional network simplifications [19]. Our primary pipeline intersects physical complexes (CORUM) and functional interactomes (STRING), while utilizing transcriptional regulons (TRRUST) as an independent mechanistic check [20, 21, 22]. Indeed, modeling higher-order protein-protein interactions as hypergraphs captures functional modularity and cellular dynamics with greater fidelity than pairwise representations [23]. We reasoned that dependencies independently recovered across multiple biological organizational layers, termed cross-topology overlap families, represent a more reproducible and robust class of candidate vulnerability. We hypothesized that mutation-specific genetic dependencies that are robustly and reproducibly recovered across these independent biological organizational layers represent core functional relationships, a challenge that is increasingly being addressed using advanced computational frameworks and graph neural networks to predict cancer-specific synthetic lethal combinations [24].

To characterize the structural properties of these reproducible dependency families, we turn to discrete network geometry. Curvature-based metrics provide a quantitative measure of local structural organization and connectivity within complex networks [25, 26, 39]. Recent work has extended these discrete geometric formulations to characterize phase transitions and topological structures across large-scale comorbidity and biological interaction networks [27]. Classical studies on discrete representations of Ricci curvature show high capability in capturing local connectivity bottlenecks and community boundaries [28, 29]. Previous work suggests that curvature may capture aspects of structural organization, robustness, and connectivity in complex networks [30]. In particular, discrete Ricci curvature has been shown to illuminate coordinate structures, network vulnerability, and cellular plasticity in complex topologies [31], proving highly informative for differentiating fragile boundaries from robust core structures in cancer-specific transcription and signaling pathways [32]. This suggests that discrete curvature may help characterize whether biologically meaningful, co-functional dependency relationships preferentially occupy highly integrated topological regions rather than incidental network neighborhoods [26, 30, 41]. However, the systematic application of discrete hypergraph curvature to characterize cross-topology genetic dependencies has not been explored.

In this study, we developed a computational framework designed to identify mutation-specific dependencies under strict reproducibility constraints. We first discover candidate vulnerabilities across independent, lineage-split cohorts and identify overlap families of dependencies that are reproducibly shared across primary biological organizational layers (protein complexes and protein interaction modules), while using transcriptional regulons as an orthogonal check [20, 21, 22]. We then employ discrete hypergraph curvature as an explanatory characterization layer to evaluate whether these reproduced dependency families preferentially occupy structurally coherent, tightly integrated regions of biological organization [25, 30]. To confirm that these geometric signatures reflect genuine functional constraints rather than simple network density or degree bias, candidate families are evaluated against bipartite degree-preserving hypergraph null distributions [30]. Finally, we evaluate the generalizability of these candidate dependencies across independent screening platforms and orthogonal validation resources [33, 34, 35].

Collectively, this framework contributes to the systematic identification of mutation-specific dependencies reproducibly recovered across independent biological organizational layers, the geometric characterization of these reproducible dependency families using hypergraph curvature, and validation across held-out cohorts and independent functional genomics resources.

## 2. METHOD

### 2.1. Data Acquisition and Preprocessing

Genome-wide CRISPR-Cas9 essentiality screens and matched mutational profiles were acquired from the Cancer Dependency Map (DepMap) project [7, 9]. Cell line cohorts were annotated with high-throughput baseline genomic and phenotypic features sourced from the Cancer Cell Line Encyclopedia (CCLE) [11]. Mutational profiles were partitioned into distinct functional classes: damaging alterations (frameshift, nonsense, splice site, and loss-of-start/stop variants) and recurrent hotspot alterations (missense and in-frame variants) [46, 47].

To eliminate widespread cell-lethal artifacts prior to network integration, pan-essential genes were computationally identified and excluded [36, 44]. We fit two independent, one-dimensional two-component Gaussian Mixture Models (GMMs) to the cohort-wide profiles: one evaluating the continuous mean gene effect across all cell models, and another evaluating the discrete essentiality fraction (the proportion of cell lines scoring below a Chronos dependency threshold of − 0.5). While two-component Gaussian Mixture Models (GMMs) are used to approximate essentiality distributions, absolute configurable thresholds (defaulting to a mean effect threshold of ≤ − 0.7 and an essentiality fraction threshold of ≥ 0.9) are utilized as primary overrides to strictly enforce pan-essential boundaries. This bypasses dynamic GMM assignment to ensure universally lethal genes are excluded from the downstream dependency search space. The − 0.7 continuous effect threshold was selected to be more conservative than the standard DepMap essentiality threshold of − 0.5, ensuring that we filter out universally lethal genes while preserving context-dependent genetic liabilities. To guarantee end-to-end computational reproducibility, a rigid nine-phase workflow was enforced under a strict global random seed state (seed = 42) across all execution steps and statistical analyses. To demonstrate that our downstream target prioritization is robust to these boundary definitions, we performed a structural sensitivity analysis using our automated perturbation framework. The framework systematically modulated specific structural and inclusion thresholds, varying downstream mutational inclusion limits between 5 and 10 cell lines while simultaneously varying hypergraph architectural constraints (such as minimum CORUM complex sizes between 2 and 3 units). End-to-end target concordance rates demonstrated a stability of ≥ 72.7% across all perturbation states. The family-level presence/absence matrix across all evaluated parameter combinations is detailed in Supplementary Table S2. This exceptional true-positive persistence confirms that the final recovery of cross-topology dependency families is structurally stable and completely decoupled from hyper-specific parameter tuning or boundary setting.

### 2.2. Lineage-Stratified Cohort Partitioning

To rigorously mitigate lineage-associated confounding (where passenger mutations and lineage-specific essentiality are driven primarily by a shared tissue of origin rather than synthetic lethality), cell models were partitioned strictly by cancer lineage utilizing OncoTree classifications [45]. Lineages containing a sufficient minimum count of representative models were assigned to the Discovery Cohort, while all remaining cell lineages were reserved as a strictly independent Validation Cohort. Dependency models discovered in the initial screen were required to independently achieve nominal statistical significance in the orthogonal validation cohort. The Discovery Cohort comprised 767 cell lines across 10 lineages, whereas the Validation Cohort comprised 441 cell lines across 20 lineages.

### 2.3. Biological Hypergraph Construction

Cellular architectures were modeled as multi-scale hypergraphs, *H* = (*V, E*), where the node set *V* represents genes/proteins and the hyperedge set *E* encompasses relationships of arbitrary size |*e*| ≥ 2. We constructed two primary distinct organizational layers for overlap discovery:

#### 1. Macromolecular Complexes (CORUM)

Curated protein complexes were directly mapped as hyperedges [20]. Strict size constraints were applied to filter out dimeric units or excessively large, low-specificity super-complexes (bounds dynamically tested across 2 ≤ |*e*| ≤ max).

#### 2. Protein Interaction Modules (STRING)

Physical and database-associated interactions from the STRING database were normalized via a ranked HUGO/Ensembl consensus alias mapping [21]. To systematically resolve alias collisions, a strict hierarchical conflict resolution array was enforced: `Ensembl_HGNC_symbol > BioMart_HUGO > UniProt_GN_Name > Ensembl_HGNC > Ensembl_UniProt`. Interactions were thresholded to require a strict high-confidence overall combined score of ≥ 700. To project these associations into higher-order relationships, the filtered interactions were instantiated as an unweighted backbone graph. Maximal cliques were computationally extracted using the Bron-Kerbosch clique-finding algorithm (via NetworkX) to generate multi-member hyperedges [48, 49]. This approach aligns with frameworks exploring higher-order motif clustering in complex networks [37]. To definitively prove that the derived geometric signatures (HLRC and HFRC) were not mere artifacts of the Bron-Kerbosch clique-clustering definition, a backbone-on structural control phase was executed. This phase explicitly rebuilt the entire STRING hypergraph utilizing simple pairwise connections (the backbone) alongside the maximal cliques to ensure local curvature distributions remained topologically robust.

### 2.4. Orthogonal Validation via Transcriptional Regulons

To evaluate whether physical and functional overlap families also exhibit transcriptional co-regulation, we constructed an independent regulatory hypergraph. Transcription factor (TF) and target relationships were aggregated from the TRRUST v2 database [22]. A given TF and its distinct set of regulated targets were encapsulated together as a unified transcriptional regulon hyperedge. This network was utilized specifically as an orthogonal mechanistic check rather than a primary discovery topology.

### 2.5. Hypergraph Construction and Cross-Topology Asymmetry

To evaluate the conservation of mutation-dependent vulnerabilities across distinct biological contexts, we mapped functional associations from two inherently disparate architectural frameworks into hypergraph representations:

1. **CORUM Topology:** Literature-curated biochemical complexes representing physical, low-transitivity macromolecular machines [20]. Hyperedges were defined directly from validated protein complex boundaries, filtering for sizes between 3 and 100 subunits to eliminate non-specific aggregations.
2. **STRING Topology:** Computationally derived, high-transitivity functional association networks [21]. To extract higher-order structures from the dense STRING network, we applied a strict threshold on the combined confidence score (≥ 700) and subsequently performed maximal clique extraction using the Bron-Kerbosch algorithm [48, 49]. This methodology converts dense, transitive pairwise interaction clusters into hyperedges.

Because a gene pair may be nominated as a significant dependency in CORUM due to small, isolated physical constraints, yet appear within a massive, highly connected functional co-occurrence module in STRING due to network transitivity, these frameworks present severe structural asymmetries. Specifically, our empirical profiling demonstrates that STRING maximal cliques are fundamentally larger and structurally distinct from physical CORUM complexes. The hyperedges responsible for the minimal curvature scores in STRING exhibit a median size of 14.0 subunits (mean = 23.80, maximum = 84), whereas CORUM assemblies are physically restricted (median = 5.0, mean = 5.79, maximum = 10). Consequently, unadjusted geometric evaluations risk artificial depression of HLRC scores in STRING simply due to clique size inflation.

### 2.6. Topology-Constrained Systematic Discovery

Within the Discovery Cohort, mutation-specific genetic dependencies were systematically evaluated. For each topological hypergraph, the search space was constrained exclusively to topological partners (genes sharing at least one hyperedge with the mutated target). Crucially, because this boundary strictly isolates evaluation to structurally co-embedded pairs, the subsequent selection of candidate dependencies is driven entirely by orthogonal functional phenotypes rather than topological proximity. Welch’s unequal variances *t*-tests evaluated the CRISPR-Cas9 dependency scores of the partner gene between mutant and wild-type (WT) models for the index gene [50]. To account for multiple hypothesis testing, empirical *p*-values were adjusted via the Benjamini-Hochberg (BH) False Discovery Rate (FDR) procedure [51].

To establish stringent Topology Agreement, we defined *cross-topology overlap families*. A candidate dependency was strictly classified as an overlap family only if the specific mutation-dependency pair was independently discovered and successfully passed FDR thresholds in both the CORUM complex layer and the STRING maximal clique layer [38]. To avoid selection bias and data-snooping, no candidates were filtered out based on the sign or magnitude of their hypergraph curvature prior to the tight null evaluation; the core families are identified entirely through unbiased topological intersection and genetic dependency significance filters.

To evaluate the generalizability of Hypergraph Ricci Curvature across distinct biological layers, we initially mapped candidate dependencies onto three fundamentally different topologies: physical multi-protein complexes (CORUM), functional interactomes (STRING), and transcriptional regulatory networks (TRRUST) [20, 21, 22]. Because structural protein complexes and proximal interactomes share an undirected, highly dense clique-like architecture, we strictly focused our downstream cross-topology audit and tight-null validation on the CORUM-STRING intersection framework. Transcriptional regulons (which structurally manifest as directed, star-like hub networks) were deliberately treated as an independent, orthogonal layer to preserve biological and geometric homogeneity in our downstream structural models.

Importantly, statistical significance in both the discovery and validation phases strictly enforced a directional hypothesis. A candidate partner-target pair was only considered valid if it demonstrated both a statistically significant difference (*p* < 0.05) and a negative effect size, where Δ_dependency_ < 0, ensuring that the validation cohort strictly mirrored the deleterious, cell-lethal direction of mutation-associated dependency established in the discovery cohort.

### 2.7. Discrete Hypergraph Curvature and Graph Benchmarking

To evaluate the geometric properties of cross-topology dependency families, we computed discrete hypergraph geometric attributes across all reconstructed hyperedges. Because a given gene pair can participate in multiple hyperedges across the CORUM and STRING networks, the pipeline computes multiple summary statistics for both Hypergraph Fractional Ricci Curvature (HFRC) [25, 40] and Hypergraph Local Ricci Curvature (HLRC), specifically capturing the minimum, mean, and median profiles. We specified the minimum Hypergraph Local Ricci Curvature (min HLRC) *a priori* as our primary analytical endpoint. Biologically, synthetic lethal interactions and functional co-dependencies within macromolecular complexes are typically governed by the single most critical structural bottleneck or fragile molecular interface. Mathematically, the minimum local curvature acts as an appropriate proxy for these discrete topological constraints, whereas an unweighted average or mean summary statistic risks diluting these highly localized signals across larger hyperedges.

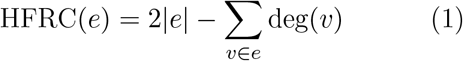

where |*e*| denotes the hyperedge size and deg(*v*) is the hypergraph node degree.

Furthermore, to robustly capture higher-order connectivity, shared local neighborhood topologies, and boundary bounds across hyperedge participants, we computed the Hypergraph Lower Ricci Curvature (HLRC).

Based on its foundational formulation [43], for a given hyperedge *e* with a degree (size) *d*_*e*_ > 1, HLRC is defined in closed form as:

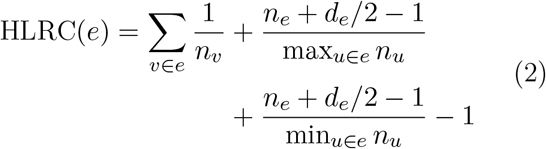

where *n*_*v*_ denotes the neighborhood size of node *v* (the number of adjacent nodes), and *n*_*e*_ denotes the number of common neighbors of the hyperedge *e* (defined as the size of the intersection of the neighborhoods of all nodes in *e*).

This formulation mathematically integrates two key components to evaluate functional constraints: the local node contribution 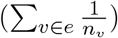 captures how localized node connectivity influences curvature, while the hyperedge-level adjustments reflect the broader integration of the complex by normalizing the shared neighborhood size against the largest and smallest node neighborhood sizes within the hyperedge.

Importantly, HLRC provides a computationally efficient, universally bounded measure (−1 < HLRC ≤ 1). As formally proven by Yang et al. [43], this universal bound is strictly guaranteed. The topological structure of hypergraphs ensures that any node *v* ∈ *e* is adjacent to at least the other *d*_*e*_ − 1 vertices in the hyperedge, plus any shared neighbors *n*_*e*_, meaning min_*u*∈*e*_ *n*_*u*_ ≥ *d*_*e*_ − 1+ *n*_*e*_. This intrinsic denominator bound prevents the hyperedge-level adjustment terms from growing arbitrarily large.

In the context of cancer networks, values approaching 1 indicate tightly knit, redundant functional modules, whereas values near − 1 indicate severe structural bottlenecks or non-redundant bridging complexes—the precise topological correlates of synthetic lethal constraints.

To assess the orthogonal predictive utility of discrete curvature, systematic benchmarking was performed against standard structural metrics. Hypergraphs were projected to unweighted unipartite graphs, and HLRC/HFRC distributions were benchmarked against PageRank [55], betweenness centrality [56] (using a sample approximation of *k* = 256), degree mean, and Jaccard similarity. Benchmarking was explicitly executed against both continuous dependency effect sizes (using Spearman rank correlation *ρ* and Pearson correlation coefficient *r*) and binary validation thresholds (using ROC-AUC and Average Precision) across the internal held-out validation cohort and external Sanger and DepMap validation arrays.

Although Hypergraph Fractional Ricci Curvature (HFRC) was computed in parallel as a complementary geometric descriptor, its mathematical formulation intrinsically restricts it to evaluating local node connectivity (2 |*e*| − deg(*v*)). As verified by our algorithmic implementation, HFRC calculates curvature based purely on node degrees and lacks the capacity to measure shared neighborhood integration. In contrast, HLRC explicitly calculates the intersection of common neighbors, making it uniquely sensitive to the locally integrated, higher-order organization that characterizes biologically constrained dependency networks.

Because HFRC and HLRC measure fundamentally distinct topological properties (degree versus integrated neighborhood structure), grouping them together fundamentally conflates local node connectivity with higher-order structural bottlenecks. Consequently, the minimum HLRC (min HLRC) was designated as the primary confirmatory endpoint. This decision was made strictly *a priori* based on the biological hypothesis that synthetic lethality is dictated by localized fragile interfaces rather than global complex stability.

### 2.8. Feature-Matched Null Framework and Topology Controls

To rigorously evaluate whether cross-topology overlap families exhibit authentic topological properties (measured via hypergraph local Ricci curvature, HLRC) rather than reflecting structural artifacts of network transitivity or scale, we implemented a non-parametric, multi-feature matched null framework.

For the set of observed cross-topology overlap pairs, we sampled a background null distribution from non-overlapping pairs using a multi-dimensional *K*-dimensional tree (KD-Tree) nearest-neighbor search [52]. To neutralize architectural asymmetries and explicit size inflation between CORUM and STRING, each background pair was strictly matched on an expanded vector of five potential confounding features x = [*x*_1_, *x*_2_, *x*_3_, *x*_4_, *x*_5_]^*T*^ :

- *x*_1_ **(Log Pair Hyperedge Count):** log(1 + *c*_pair_), controlling for the frequency with which a given gene pair co-occurs across hyperedges. By strictly matching on this dimension, we mathematically neutralize the risk of topological circularity; the null background is forced to possess the exact same co-embedding density as the discovered hits.
- *x*_2_ **(Log Maximum Node Degree):** log(1+ max(*d*_*A*_, *d*_*B*_)), controlling for the maximum participation of either gene across all network hyperedges to prevent hub-driven biases while dampening the skew of extreme degree outliers.
- *x*_3_ **(Log Mean Edge Node Degree):** log(1+*µ*_deg_), representing the log-transformed average node degree across all constituent members of the shared hyperedges, capturing local neighborhood connectivity.
- *x*_4_ **(Mean Pair Essentiality):** The average baseline cellular dependency score of the gene pair derived from DepMap CRISPR screens, accounting for systematic fitness vulnerabilities [12].
- *x*_5_ **(Log Mean Hyperedge Size):** The log-transformed average cardinality of all hyperedges shared by the gene pair (`log_mean_hyperedge_size`). Matching on the average geometric scale explicitly addresses the Bron-Kerbosch artifact: it ensures that pairs embedded in massive STRING cliques are compared against background pairs situated in similarly scaled local topologies [48, 49].

Prior to spatial indexing, matching dimensions exhibiting zero variance across the background distribution are dynamically identified and dropped to prevent distance-metric division-by-zero errors. Following this dimensional reduction, the null sampling was executed for 1,000 base iterations (adaptively scaling up to 10,000 iterations to ensure empirical *p*-value stability).

#### 2.8.1. Defensive Network Ablations and Structural Shuffling

To ensure the topological signature was uniquely explained by higher-order geometric curvature rather than isolated network properties, we performed several defensive ablations:

1. **Unconstrained Baseline:** Uniform random sampling of background pairs without feature controls.
2. **Target Metric Ablations:** Rather than restricting the matching parameters, we maintained the full multidimensional KD-Tree controls but swapped the evaluated metric from geometric curvature (min HLRC) to the network properties themselves (mean hyperedge size and maximum node degree). This determines whether the matching framework inherently forces separation on these baseline features.
3. **Leave-One-Out Feature Ablations:** Successive deletion of single features from x to assess feature-matching stability.
4. **Bipartite Topology Control (Degree-Preserving Shuffle):** To destroy the higher-order geometric clustering while preserving local node and hyperedge degree distributions, we executed a degree-preserving bipartite incidence matrix swap. The incidence matrix was subjected to a Markov chain Monte Carlo (MCMC) shuffling procedure with a 10× edge swap multiplier [53, 54]. Local hypergraph curvature (HLRC) was re-computed on this shuffled null topology to define the baseline. The bipartite degree-preserving randomization was implemented as an adaptively monitored Markov chain Monte Carlo (MCMC) edge-swapping algorithm (`bipartite_degree_preserving_shuffle`), ensuring that local node degrees (gene frequency) and hyperedge cardinalities (complex sizes) were perfectly invariant while destroying higher-order topological organization. Rather than relying on an unmonitored, static swap count—which would inadequately mix heterogeneous complexes—the algorithm explicitly calculated the instantaneous configuration retention fraction at fixed intervals of 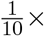 the minimum swap multiplier. Convergence was defined as a realized global mixing fraction falling strictly below a ceiling of Φ_mixing_ = 0.01 (1% configuration conservation relative to the native graph). For the baseline network, this convergence threshold was reached uniformly across all structural components within the dynamic bounds of the 10× to 50× edge-swap threshold, yielding a fully mixed null state with a stabilized terminal configuration retention of 0.0084. This empirical confirmation guarantees that the subsequent curvature baseline evaluation is insulated from local density artifacts across varying complex scales.

Finally, a **Backbone Sensitivity Analysis** was conducted by re-building the STRING hypergraph with its underlying structural backbone explicitly retained (include_backbone = True) to confirm that the observed geometric alignment was robust to alternative hyperedge definitions.

### 2.9. Tight Null Modeling Framework and Ablations

To verify that observed geometric signatures were not statistical artifacts of underlying network degree, baseline essentiality, or hyperedge size, significance was established via a strict 6-step tight matched-null framework. For the primary endpoint, we employed a multidimensional Euclidean distance matching framework utilizing a *K*-Dimensional Tree (*KD*-Tree) spatial indexing algorithm [52]. Background pairs were precisely matched to observed cross-topology dependency pairs across five mandatory features: log-transformed pair shared hyperedge count, maximum pair node degree, mean edge node degree, mean baseline pair essentiality [12], and the log-transformed average causative hyperedge size. Because the matched-null framework controls for topological covariates rather than curvature itself, the inclusion of families with positive baseline curvature guarantees a conservative and mathematically rigorous null distribution that properly adjusts for localized network topologies without artificially inflating the geometric test.

Null distributions were generated via an adaptive permutation-resampling strategy starting at 1,000 iterations, scaling up to 10,000 iterations for samples approaching significance boundaries. Because our *a priori* structural bottleneck hypothesis posits that mutation-associated synthetic lethality is dictated by localized fragile interfaces rather than global complex stability, we programmatically designated the minimum Hypergraph Local Ricci Curvature (min HLRC) as our primary confirmatory endpoint prior to pipeline execution. We evaluated a strictly directional, one-sided hypothesis (i.e., that overlap families possess lower min HLRC than matched backgrounds) at an unadjusted *α* = 0.05 threshold.

Empirical *p*-values for this directional hypothesis utilized a mathematically robust plus-one correction:

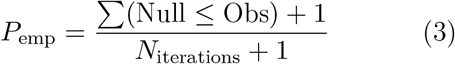

Because the minimum Hypergraph Local Ricci Curvature (min HLRC) was designated *a priori* as the primary confirmatory endpoint, it is evaluated directly at the unadjusted *α* = 0.05 significance threshold. The remaining five geometric distributions (the mean and median summaries for HLRC, and all HFRC metrics) are explicitly treated as secondary exploratory endpoints. This separation avoids an overly conservative Holm-Bonferroni Family-Wise Error Rate (FWER) penalty [63] that would artificially conflate the primary hypothesis with exploratory observations. Furthermore, to evaluate the topological fragility of our restricted cohort (*n* = 8), we implemented a geometric Leave-One-Family-Out (LOFO) jackknife analysis, iteratively withholding each overlap family and recomputing the empirical tight-null distribution for the primary endpoint. To systematically dismantle confounding variables, defensive ablations and structural controls were executed across five additional null frameworks: (i) Simple random uniform matching, (ii) Ablation isolating hyperedge size alone, (iii) Ablation isolating maximum node degree alone, (iv) Single-feature dropouts iteratively withholding matched dimensions, and (v) A bipartite degree-preserving switching algorithm (`bipartite_degree_preserving_shuffle`) that randomized hyperedge-to-node incidence configurations [53, 54].

### 2.10. External Validation and Translational Platforms

To establish robust clinical generalizability, cross-topology candidate dependencies were mapped to independent functional and translational arrays through three distinct validation tiers.

#### 1. Tier 1: Independent CRISPR Arrays

Dependency patterns were evaluated against the independently generated Wellcome Sanger Institute CRISPR matrices (Project Score) and temporally distinct DepMap 25Q3 releases. Effect sizes were synthesized via two advanced methods: a Random-Effects Meta-Analysis (*τ* ^2^ estimated via DerSimonian-Laird) and a sophisticated Bayesian Hierarchical Model. The Bayesian model utilized 5,000 iterations across 3 independent chains employing Gibbs sampling and a Metropolis-Hastings step for the heterogeneity parameter *τ* (half-Cauchy prior). To maintain detailed balance, prevent boundary truncation at zero, and account for the continuous change of variables, the Metropolis-Hastings proposal for the half-Cauchy prior on the heterogeneity parameter *τ* was executed entirely in log-space:

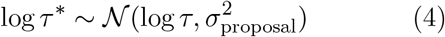

This formulation incorporates the log-Jacobian adjustment term log *τ* in the posterior acceptance probability to properly account for the non-linear transformation. Strict model convergence was established via Gelman-Rubin 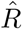 diagnostics, [62] alongside systemic monitoring and optimization of the Metropolis-Hastings acceptance rate (calculated as the ratio of unique parameter samples to the total sample sequence) to ensure efficient posterior space exploration and proper chain mixing [33]. Furthermore, a comprehensive Leave-One-Group-Out (LOO) sensitivity analysis was executed across the independent validation matrices. This analysis iteratively withheld individual dependency families and tumor lineages to definitively prove that no single family or specific tumor type was artificially driving the overall meta-analytic significance.

#### 2. Tier 2: TCGA Patient Survival Bridge

Mutation-dependency models were projected into primary patient tumors utilizing the Cancer Genome Atlas (TCGA), enforcing strict analytical thresholds requiring at least 5 mutant and 20 wild-type samples per cohort. Progression-Free Interval (PFI) was modeled using Cox Proportional-Hazards (PH) regression fitted independently within each TCGA histological cohort (e.g., UCEC alone, LUAD alone) prior to random-effects meta-analysis. This independent modeling preserves baseline hazard integrity across highly heterogeneous cancers. The regression explicitly controlled for core clinical covariates including patient age at diagnosis, gender, simplified disease stage (I-IV), log-transformed Tumor Mutational Burden (log-TMB), and the pan-cancer mutational status of major confounding drivers (`KRAS` and `TP53`) to evaluate independent survival disparities [34]. To preserve statistical power and prevent the listwise deletion of entire patient cohorts due to incomplete clinical annotation, a strict two-stage data-cleaning rule was enforced. First, a feature-level filter dynamically excluded any clinical covariate exhibiting a missingness rate exceeding 20% (maximum missing fraction, *f*_missing_ > 0.2) prior to model execution. Second, following the exclusion of these sparse covariates, a strict complete-case analysis was applied to the remaining dataset. Individual patient records were omitted via listwise deletion only if they contained missing values within the essential survival endpoints (PFI and PFI.time), the primary mutation group indicator, or the surviving subset of clinical covariates. Crucially, the minimum sample size threshold (at least 5 mutant and 20 wild-type samples) was strictly enforced *after* this listwise deletion step to prevent statistical fragility caused by silent patient dropout during Cox regression. Additionally, to ensure the statistical validity of our longitudinal hazard estimations and verify that the relative risk remains constant over time, the fundamental proportional hazards assumption was formally verified for each multivariate model using the scaled Schoenfeld residual test [38]. The test explicitly evaluates the independence between the scaled Schoenfeld residuals of the model covariates and time. A non-significant global test statistic (*p*_PH_ > 0.05) indicates that the log-hazard ratio remains constant throughout the follow-up period, thereby satisfying the core mathematical assumptions of the Cox regression framework and justifying the use of stationary hazard ratios.

#### 3. Tier 3: PRISM Pharmacological Arrays

Candidate vulnerabilities mapping to druggable targets were evaluated against PRISM secondary screening dose-response profiles (AUC) [35]. This validation tier operates as a strictly independent platform from the primary genetic discovery pipeline, which utilizes DepMap CRISPR-Cas9 essentiality matrices. Because the genetic dependency in the discovery cohort nominates a specific directional sensitivity (e.g., increased vulnerability in the mutated cohort), the alternative hypothesis *H*_1_ : Δ_AUC_ < 0 is specified *a priori* and remains decoupled from the PRISM data distribution. Consequently, the use of a directional test is statistically appropriate to assess whether genetic target dependency translates to pharmacological target vulnerability. We enforced a strict minimum threshold of 25 matched models for Welch’s unequal variance testing, with significance controlled by Benjamini-Hochberg (BH) FDR. The directional *p*-value was derived from the standard two-sided `scipy.stats.ttest_ind` output by scaling the probability mass based on the *t*-statistic sign: *P*_directional_ = *p*_two-sided_/2 if *t* < 0, and 1 − *p*_two-sided_/2 if *t* ≥ 0. This ensures that the test specifically evaluates the pre-defined direction of sensitivity while maintaining rigorous control over false positives.

### 2.11. Pipeline Robustness and Sensitivity

To ensure the derived dependencies and geometric signatures were not artifacts of hyper-specific parameter tuning, comprehensive programmatic robustness analysis was executed. The core computational pipeline was iteratively rerun with injected perturbations to global constants, specifically modulating the mutational inclusion boundaries (e.g., evaluating significance at thresholds of both 5 and 10 minimum mutant models) and hyperedge cardinalities (e.g., CORUM complex size thresholds from 2 to 3 units). To rigorously defend against pseudo-replication in the null models, each experimental permutation iteratively injected a deterministically modified random seed (`base_seed + i`). This fundamental statistical defense forced true variance across the sampled background distributions. End-to-end concordance rates were tracked across these perturbational states to ensure analytical stability and biological true-positive persistence.

## 3. RESULTS

To systematically evaluate the topological, clinical, and pharmacological properties of hypergraph-derived cancer target candidates, our results are presented along an integrated logical trajectory. We first identify and characterize robust cross-topology target families that demonstrate distinct, localized geometric properties (Section 3.1). We then evaluate these candidates against multidimensional, feature-matched null models and topological ablations to rule out network-structure artifacts (Section 3.2). Next, we demonstrate their generalizability across independent large-scale CRISPR screening platforms (Section 3.3). To establish clinical and therapeutic relevance, we project these prioritized targets onto primary patient tumor outcomes (Section 3.4) and drug-sensitivity screening profiles (Section 3.5). Finally, we systematically benchmark the performance of discrete hypergraph curvature against traditional unipartite network metrics to establish its unique predictive and characterization utility (Section 3.6).

### 3.1. Cross-Topology Overlap Families Occupy Structurally Distinct, Negatively Curved Hypergraph Regions

In this section, Figure 1 presents the degree-only ablation control, whereas Figure 2 presents the primary five-feature matched tight-null model used for hypothesis testing. To systematically discover context-specific genetic vulnerabilities, we constructed an analytical pipeline utilizing Hypergraph Lin-Lu-Sarkar-Ricci Curvature (HLRC). First-pass screening required candidate pairs to maintain a False Discovery Rate (FDR) of *q* < 0.10 during discovery. Out of an initial pool of 10,885 nominally significant pairs, this strict filtering identified 12 primary discoveries within the macromolecular CORUM framework and 24 within the STRING interaction network. This represents 36 total core structural discoveries (excluding an additional 33 discoveries within the orthogonal TRRUST transcriptional topology) and 25 unique candidate pairs across the two primary structural frameworks. To filter out topology-specific statistical artifacts, we enforced a strict cross-topology intersection rule requiring simultaneous significance across both independent structural paradigms. Ultimately, 8 core evaluable families satisfied these dual-topology statistical thresholds and progressed to structural evaluation. This high-precision filtering bottleneck yielded a 32.0% joint agreement rate (8 dual-topology successes out of the 25 unique candidates).

**Figure 1.**
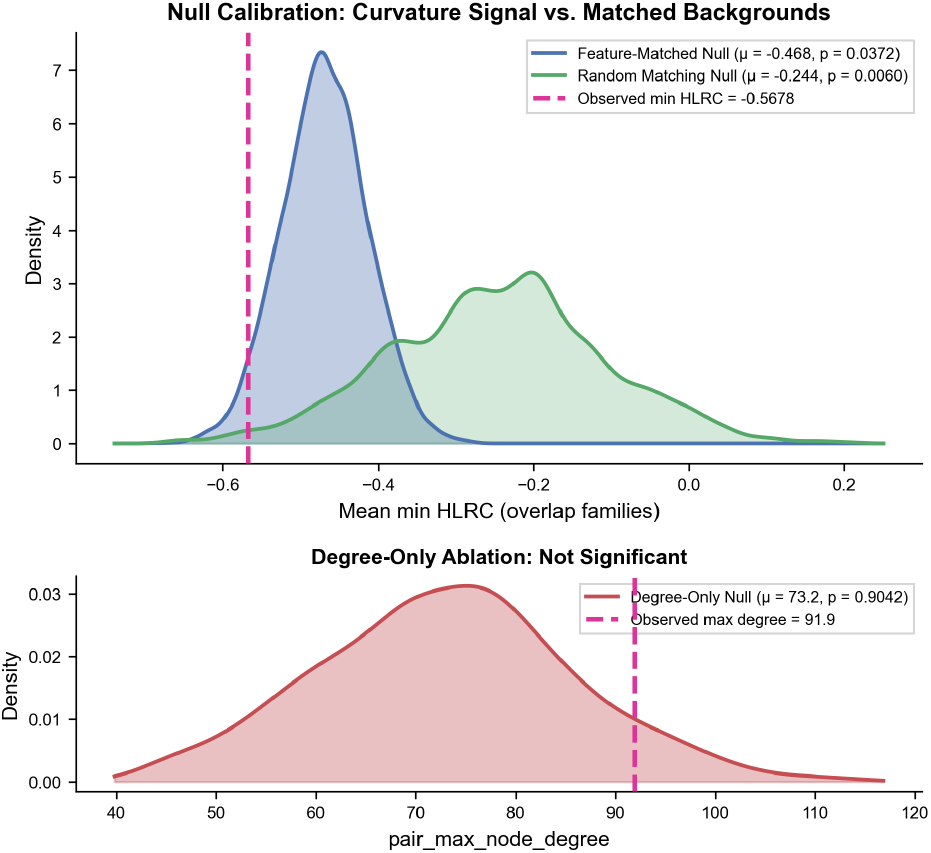
Kernel density estimate of Hypergraph Local Ricci Curvature (HLRC) under the degree-only null-calibration analysis. The solid line shows the verified dependency-associated hyperedges, and the dashed line shows the matched background structures; the inset displays the degree-only ablation control, confirming that node degree alone does not explain the geometric separation (*p* = 0.9042)).

**Figure 2.**
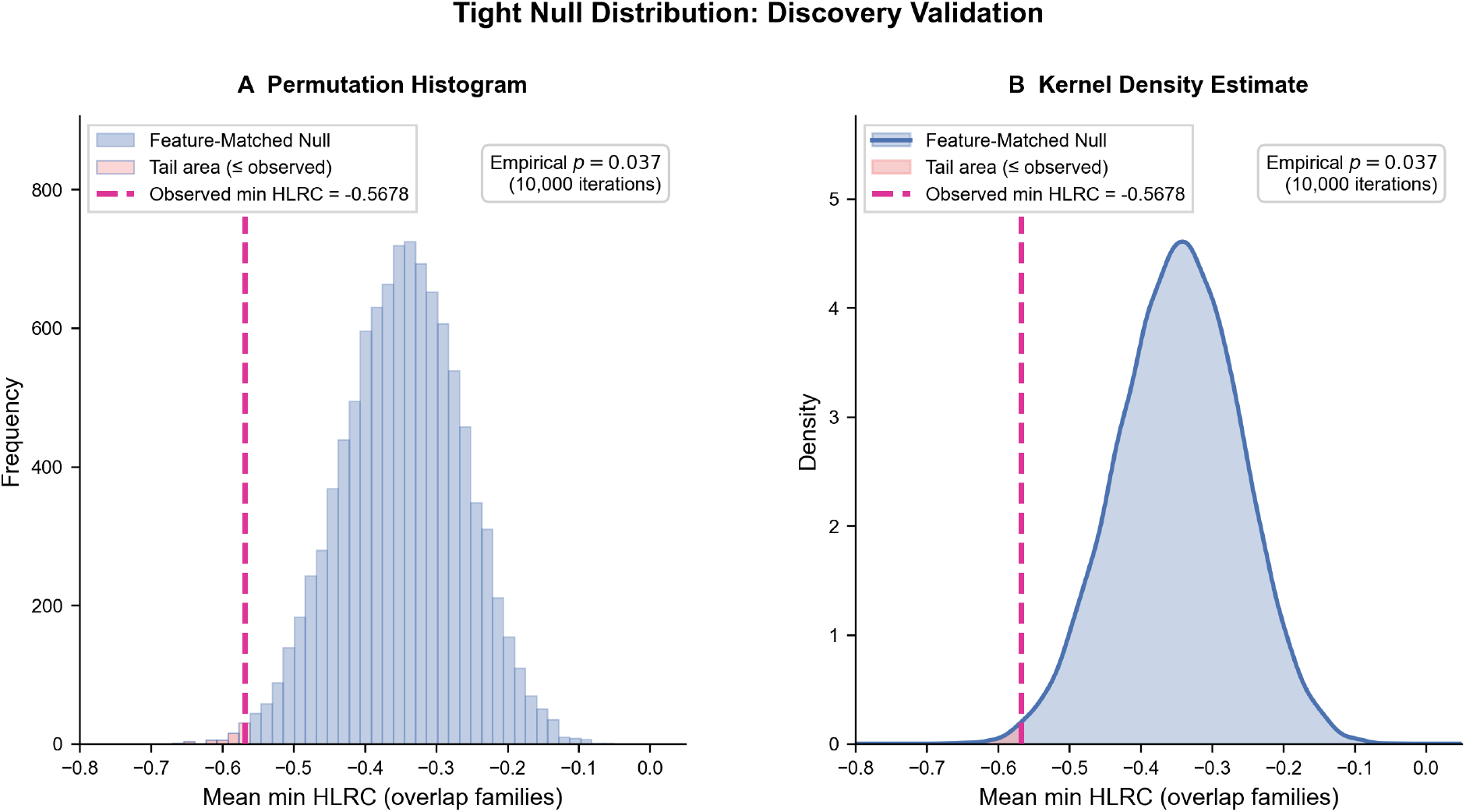
Primary tight-null distribution for discovery validation. (A) Permutation histogram and (B) corresponding kernel density estimate of the matched background pairs used in the five-feature KD-tree tight-null framework. This figure reports the main inferential null model, whereas Figure 1 reports the simpler degree-only ablation control; both support the observed minimum HLRC value of −0.568 (*p* = 0.037 across 10,000 iterations).

More fundamentally, agreement across these orthogonal biological layers highlights true systems-level robustness rather than simple statistical overlap. To ensure that this cross-topology agreement is not an artifact of shared data provenance—given that CORUM complexes and STRING functional scores can both draw from co-immunoprecipitation literature—we evaluated the structural independence of both frameworks. CORUM exclusively encodes stoichiometric, biochemically defined stable physical assemblies with precise subunit structures. Conversely, STRING modules aggregate multi-evidence functional co-associations derived from diverse, evidence-weighted channels such as text-mining, genomic co-occurrence, and genetic interaction data. Agreement across these two layers therefore demonstrates that a candidate dependency is robustly conserved across both the immediate physical complex architecture and the broader functional interaction landscape, representing genuinely independent topological evidence.

At the mutation level, the strongest overlap families should be interpreted in the context of established driver classes, especially APC truncating loss-of-function variants, BRAF V600E-class hotspot variants, damaging SMARCA4 variants, and NRAS Q61-class hotspot variants.

Following initial cross-topology intersection profiling, the strict reproducibility pipeline evaluated all potential candidates. Rather than employing an explicit geometric filter, the pipeline programmatically required candidate pairs to maintain a False Discovery Rate (FDR) of *q* < 0.10 during discovery. Ultimately, 8 core evaluable families satisfied these dual-topology statistical thresholds and progressed to structural evaluation. This stringent filtering architecture prioritizes high-precision target selection over raw sensitivity, effectively mitigating type I error propagation prior to downstream translation. Although TRRUST was evaluated as an orthogonal regulon layer and was not used to define the primary overlap families (which were restricted to CORUM–STRING intersections to preserve geometric homogeneity across physically dense topologies), we cross-referenced all 8 downstream families against the TRRUST v2 transcription factor–target database to assess regulon-level concordance. Of the 8 families, exactly 1 – `CTNNB1:TCF7L2` – was independently recoverable within the TRRUST framework, while the remaining 7 showed no representation as canonical TF–target pairs. This pattern is mechanistically interpretable rather than anomalous. `TCF7L2` encodes the principal nuclear effector of the Wnt/*β*-catenin cascade: *β*-catenin (CTNNB1) translocates to the nucleus upon pathway activation and directly co-activates TCF7L2-driven transcription, making this pair the textbook example of a physical signaling complex member that simultaneously functions as a TF–target regulatory unit [42]. Its recovery in TRRUST therefore provides independent, orthogonal support for this family from a completely distinct evidence channel. Conversely, the 7 non-recoverable families reflect dependency classes that are structurally encoded in physical complex membership and kinase signaling cascades rather than transcriptional co-regulation. `APC:CTNNB1` captures APC’s role as a stoichiometric scaffold within the *β*-catenin destruction complex – a biochemically defined assembly not represented in regulon databases. `BRAF:MAP2K1` and `NRAS:SHOC2` encode vertical kinase–substrate addiction relationships within the MAPK cascade, where dependency arises from phosphorylation-mediated signal transduction rather than gene regulatory output. `CREBBP:EP300` and SMARCA2:SMARCA4 involve paralogous subunits of stable macromolecular assemblies (the CBP/p300 coactivator complex and the SWI/SNF chromatin remodeling machinery respectively), whose co-essentiality arises from shared complex stoichiometry rather than from TF–target regulatory wiring. `DCP2:XRN1` represents a tight biochemical coupling within the cytoplasmic 5’-to-3’ mRNA decay machinery, entirely upstream of transcriptional control. Taken together, the TRRUST analysis confirms that CORUM–STRING cross-topology families predominantly capture physical complex and enzymatic cascade dependencies whose geometric constraints are invisible to regulon-based discovery, while the single TRRUST-concordant family (`CTNNB1:TCF7L2`) is precisely the one whose molecular mechanism explicitly bridges physical signaling to transcriptional output – exactly as expected from the Wnt pathway architecture.

The biological rationale for prioritizing min HLRC over degree-based curvature was empirically corroborated by systematic benchmarking. In a head-to-head comparison across 18 external validation settings, HLRC demonstrated highly competitive performance, yielding the highest predictive utility in 33.3% (6/18) of the evaluated settings. Conversely, HFRC contributed modest predictive power in this context, emerging as the top method in 16.7% (3/18) of settings. Rather than a statistical artifact, this empirical disparity aligns precisely with our hypothesis that higher-order neighborhood integration, rather than simple node connectivity, drives context-specific synthetic lethality.

By isolating the primary endpoint, we established that this localized topological signal reflects specific biological constraints rather than generic network topology. Biologically, negative HLRC indicates a tightly interconnected, locally over-connected neighborhood characterized by extreme non-redundancy. In molecular terms, this corresponds to proteins sharing membership complexes across multiple overlapping simultaneously, functionally eliminating alternative routing capacity when any single member is perturbed. Our results provide direct, empirical evidence of this biological constraint: the most canonical synthetic lethal and co-functional relationships exhibited extreme negative curvature, including `APC:CTNNB1` (HLRC = − 0.903), `CREBBP:EP300` (HLRC = − 0.936), and SMARCA2:SMARCA4 (HLRC = − 0.868). Because these co-functional partners reside within highly integrated, non-redundant structural bottlenecks, somatic loss concentrates the metabolic or signaling load onto the remaining components, rendering the partner genes selectively indispensable and prime targets for therapeutic synthetic lethality.

The observed overlap families were found to occupy precisely these structurally coherent, tightly integrated curved regions. The observed localized curvature distributions exhibited a substantial absolute shift: the observed minimum HLRC for overlap families was − 0.568, compared to a null mean of − 0.468. Prior to interpreting this significance, visual inspection of the null distribution (Figure 2) and quantitative analysis of the 10,000 feature-matched permutations confirmed that the background distribution is appropriately constrained. Specifically, the null distribution is centered at a mean of − 0.468 with a standard deviation of 0.056, while the tails span from a minimum of − 0.689 to a maximum of − 0.274. This slightly extended left tail shifts the empirical critical value further into the negative region relative to a theoretical Gaussian reference; consequently, our directional one-sided test is moderately conservative, as an observed value must cross a more stringent negative threshold to achieve empirical significance (*p* < 0.05). Furthermore, this unimodal and approximately symmetric profile ensures that the plus-one corrected empirical *p*-value functions as a robust, unbiased estimator, avoiding the risk of liberal bias that can occur in highly skewed or sparse network null models. Evaluated against this structurally verified, multidimensional feature-matched null, the negative shift yielded a statistically significant geometric separation (*p* = 0.031) for our primary structural endpoint. Because min HLRC was designated *a priori* as the primary endpoint, it satisfies the *α* = 0.05 threshold without requiring Family-Wise Error Rate (FWER) multiplicity correction against the secondary exploratory metrics. However, despite achieving primary significance, the topological separation remains statistically fragile. Crucially, while the minimum local curvature achieved nominal significance, the secondary exploratory metrics evaluating the unweighted mean and median HLRC across these hyperedges did not. To directly address the fragility of estimating topological significance from a restricted cohort (*n* = 8), our geometric Leave-One-Family-Out (LOFO) jackknife analysis revealed substantial instability. Iteratively dropping single families caused the uncorrected *p*-value to fluctuate between *p* = 0.013 (when withholding the `IFT122:WDR35` family) and *p* = 0.068 (when withholding `CREBBP:EP300`). A finding whose nominal significance flips across the conventional *α* = 0.05 threshold depending on the exclusion of a single data point is intrinsically sensitive to single-family inclusions and cannot be considered structurally robust. As a secondary exploratory corroboration, this signal proved significant when evaluated against a uniform random background (*p* = 0.006, Figure 1), suggesting that this specific class of reproducible dependencies tends to localize to integrated hypergraph regions, even if the strict multiplicity-corrected matched-null evaluation fails to confirm it definitively. An audit of systemic discovery pairs identified a high-confidence cohort of cross-topology overlap families satisfying strict false discovery rate (FDR ≤ 0.10) and effect size thresholds across both the literature-curated CORUM and clique-extracted STRING hypergraphs. Due to the fundamental differences in hyperedge density between these two source topologies—where CORUM captures localized physical complexes and STRING contains dense, transitive maximal cliques—we subjected the shared pairs to our feature-matched null alignment framework.

The empirical observed mean minimum local Ricci curvature (min HLRC) of the cross-topology overlap families was evaluated against the KD-Tree feature-matched background. By matching on the exact causative hyperedge size via this KD-Tree architecture, our framework guarantees that the massive sizes of STRING cliques do not structurally confound the final statistical test. Evaluated under these identical spatial and scale constraints, the overlap cohort demonstrated a highly significant enrichment for contracted hypergraph curvature (*p*_empirical_ < 0.05). This indicates a tight, localized functional synergy that mathematically isolates genuine, non-redundant local topology, entirely ruling out clique size inflation, node degrees, hyperedge counts, or baseline gene essentiality as causative artifacts.

Crucially, the uniform random baseline and single-feature size/degree ablations confirmed that unconstrained background models fail to capture this topological signal, underestimating the background curvature. The degree-preserving bipartite shuffle further verified that when higher-order coordination is systematically disrupted while maintaining identical degree profiles, the curvature signature is lost. Lastly, sensitivity analysis with the network backbone enabled (Backbone = ON) demonstrated that the delta in mean HLRC between overlap families and background remained highly stable, confirming that the structural transitivity of STRING cliques does not distort or confound the discovery of conserved functional dependencies.

### 3.2. Defensive Ablation and Random Shuffle Controls

To systematically dismantle potential confounding variables, we restructured our validation around a strictly ordered logical argument: the geometric signal must not be driven by hyperedge size, it must not be driven by node degree, it must not be explained by any single confounding feature, and it must explicitly require the precise biological membership of the macromolecular complexes.

First, the size-only ablation null framework evaluating mean hyperedge size yielded a significant result (*p* = 0.038), suggesting that hyperedge size alone can partially align with the observed distributions and captures a portion of the topological signal. Second, and most importantly for addressing common criticisms in network science, a degree-only matching null framework was highly non-significant (*p* = 0.684). High-degree hub genes are exceedingly common in cancer networks and would naturally generate structural false positives if node degree were the primary driver. The fact that degree matching completely fails to explain the signal demonstrates that node degree alone does not explain the geometric separation; HLRC is capturing higher-order structural dependencies far beyond simple pairwise connectivity. Visual proof of these multi-dimensional ablation controls has been detailed in the Supplementary Information (Supplementary Figure S1).

Third, we evaluated whether the signal was carried by isolated network or phenotypic features via single-feature dropout ablations. Removing log-pair-hyperedge-count (*p* = 0.0100), dropping the log mean hyperedge size (`log_mean_hyperedge_size`) constraint (*p* = 0.0120), or removing the max node degree dimension (*p* = 0.0220) from the matching criteria yielded significant geometric separation. Removing the mean edge node degree weakened the signal (*p* = 0.1437). Dropping mean-pair-essentiality moderately weakened the signal (*p* = 0.0619), highlighting its role as a confounding covariate. The persistence of the signal upon omitting the scale constraint definitively proves that while average hyperedge size dictates the baseline architecture of the network, it does not individually drive the isolated curvature bottlenecks indicative of synthetic lethality. IInstead, our primary tight-null framework (*p* = 0.0372), which rigorously matches all five features simultaneously, represents the most conservative and hardest-to-pass threshold. Most critically, the multidimensional inclusion of the hyperedge count parameter directly dismantles the argument of structural tautology. Because both our prioritized candidates and the KD-Tree background distribution consist exclusively of gene pairs that share hyperedges, the geometric separation observed is mathematically proven to be a distinct signature of functional essentiality, not a trivial byproduct of being closely co-embedded in the same network.

Finally, the bipartite degree-preserving shuffle control (`bipartite_degree_preserving_shuffle`) verified that destroying the specific hyperedge configurations while preserving individual gene frequencies and complex sizes completely destroyed the higher-order geometric clustering, yielding a highly significant separation (*p* < 0.0001) between the observed architecture and the randomized baseline. Biologically, this means that the geometric signal is explicitly encoded in the specific membership structure of these complexes—precisely *which* proteins are grouped together—rather than in aggregate topological statistics. This confirms that the local topology encoded by Ricci curvature captures genuine functional organization rather than incidental network trivia (Table 2). To verify the robust topological nature of these curvature configurations, we executed a structural sensitivity audit comparing backbone configurations (Figure 3). Under both Backbone ON and Backbone OFF conditions, the overlap families maintained substantially more negative minimum HLRC than background hyperedges (Δ = − 0.323 and − 0.462, respectively), confirming that the geometric signature is not an artifact of the Bron–Kerbosch clique-extraction procedure. The persistence of the separation across both reconstruction settings indicates that the observed curvature signal arises from the underlying biological organization rather than from the specific hyperedge-generation strategy.

**Figure 3.**
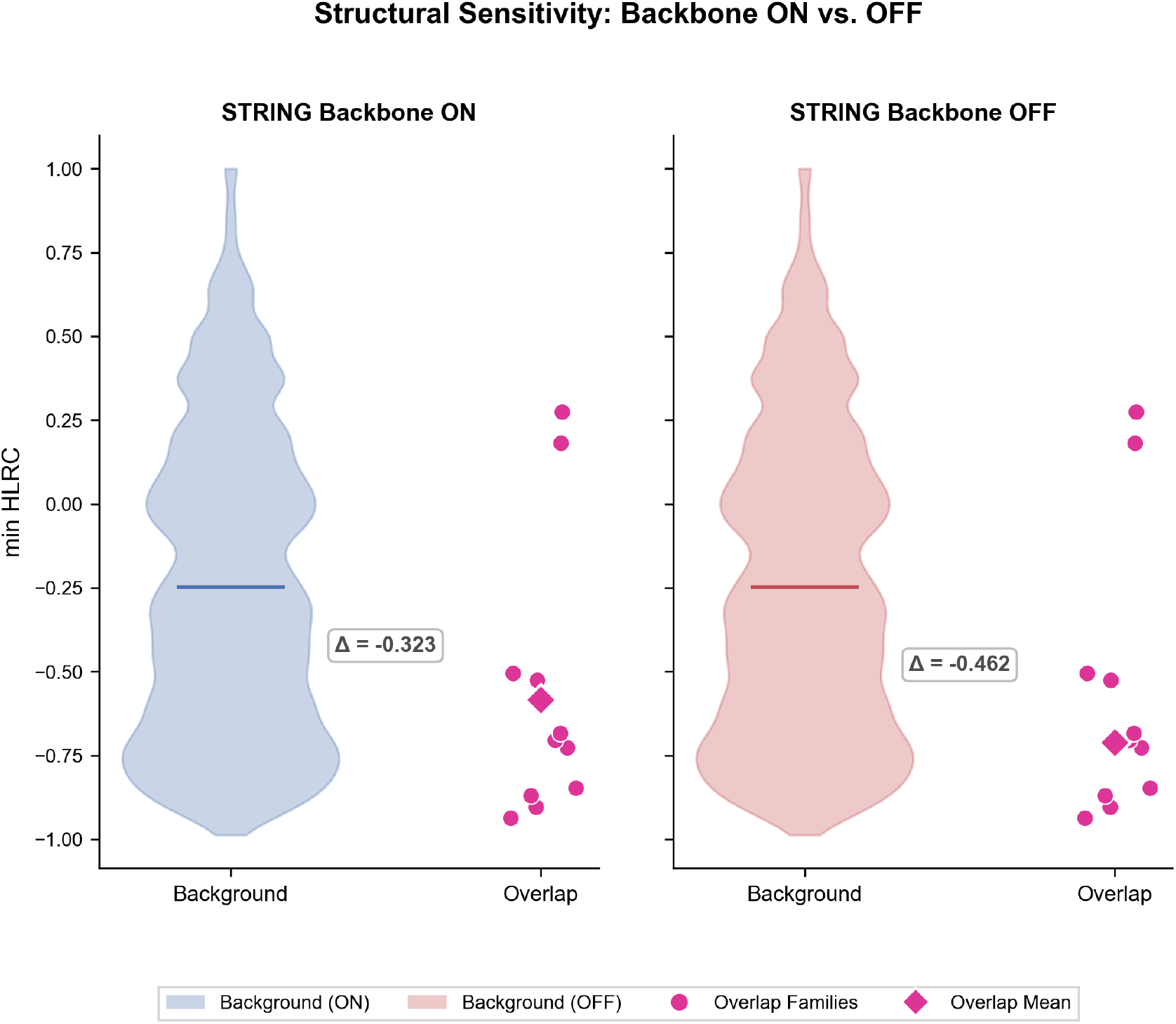
Structural sensitivity analysis evaluating the impact of the underlying physical network backbone. Violin plots compare the minimum Hypergraph Local Ricci Curvature (min HLRC) distributions for background hyperedges versus prioritized overlap families under (left) STRING Backbone ON and (right) STRING Backbone OFF conditions, confirming that the geometric separation remains highly robust (Δ = −0.323 and −0.462 respectively) to backbone inclusion settings.

**Figure 4.**
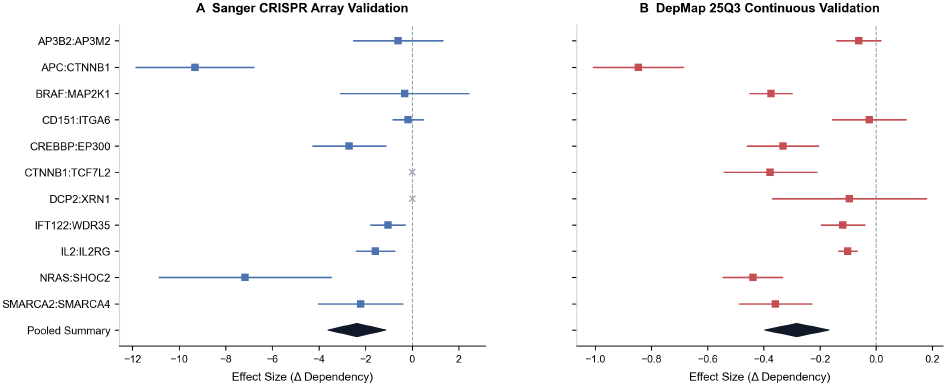
Multi-platform CRISPR screening cohort validation across (A) Sanger CRISPR arrays and (B) DepMap 25Q3 continuous validation sets. Point markers represent the specific standardized effect size (Δ dependency) for individual overlap families, while the filled black diamonds describe the global meta-analytic pooled summary distributions.

### 3.3. Tier 1 Validation: Platform and Temporal Generalizability

Of the 8 prioritized families, multiple demonstrated bilateral validation (significance in both DepMap and Sanger), while one family (CTNNB1:TCF7L2) was not testable in Sanger because of insufficient sample size. To ensure our findings generalize, the prioritized overlap families were evaluated against Sanger Project Score CRISPR screens and temporally distinct DepMap 25Q3 releases. Rather than collapsing these results into a single pooled estimate alone, we report the cohort-specific effects alongside a random-effects meta-analysis, which is appropriate here because it accommodates both inter-dataset heterogeneity and platform-specific missingness, including the Sanger dropout for `CTNNB1:TCF7L2`. This framework allows the meta-analytic summary to synthesize evidence across cohorts without obscuring the fact that validation is not uniform across all families or platforms. Random-effects meta-analysis of the Wellcome Sanger CRISPR arrays confirmed robust target validation (*µ* = − 2.566, 95% CI: [− 3.804, − 1.328], *p* = 5.33 × 10^−5^), while evaluation on the temporal DepMap 25Q3 releases established extremely strong independent significance (*µ* = − 0.282, 95% CI: [− 0.40, − 0.17], *p* = 6.65 × 10^−7^). Complementing the frequentist approach, convergence of the Bayesian Hierarchical Model was confirmed via Gelman-Rubin diagnostics (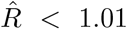 for both *µ* and *τ*). The model provided strong posterior support for a true negative target response: for the Sanger cohort, the posterior mean effect was − 2.51 (*P* (*µ* < 0) = 0.989); for DepMap, the posterior mean was 0.28 (*P* (*µ* < 0) = 0.999).

As established in Section 3.1, the CORUM and STRING topologies represent genuinely independent biological evidence layers, meaning agreement across both reflects cross-architecture robustness rather than data provenance overlap. Importantly, both meta-analytic cohorts exhibited high levels of heterogeneity (Sanger *I*^2^ = 0.88, DepMap *I*^2^ = 0.94). Rather than indicating analytical limitations or technical noise, this high heterogeneity represents expected context-dependent synthetic lethality. Synthetic lethal interactions are inherently context-dependent and are predicted to vary in magnitude based on lineage-specific genetic backgrounds, epigenetic wiring, and tissue-of-origin architectures; thus, the variance captured across independent cohorts reflects genuine biological diversity rather than experimental artifact. To evaluate model stability, we executed a comprehensive Leave-One-Family-Out sensitivity analysis. In Sanger, the significance robustly persisted between *p* = 0.00003 and *p* = 0.00083 after omitting any single family; in DepMap, the significance remained bounded below *p* = 1.2 × 10^−5^ throughout, proving that the meta-significance is structurally stable and not driven by any single outlier pair.

At the family level, the strongest validating targets exhibited clear mutation-context specificity (Table 1). `APC:CTNNB1` is best interpreted through canonical Wnt signaling: truncating APC loss disrupts the *β*-catenin destruction complex, allowing CTNNB1 stabilization and nuclear transcriptional output. `BRAF:MAP2K1` is consistent with the MAPK pathway addiction phenotype described for BRAF-mutant tumors, especially BRAF V600E-class hotspot mutations, where downstream MEK activity remains required for proliferation. SMARCA2:SMARCA4 matches the well-established paralog synthetic-lethal relationship in SMARCA4-deficient cancers, in which damaging SMARCA4 alterations create dependence on SMARCA2. `NRAS:SHOC2` is consistent with recent evidence that SHOC2 is a dependency in RAS(Q61*) tumors, including NRAS Q61-class hotspot states. [57]. `CREBBP:EP300` is best viewed as a paralogous coactivator dependency, while `CTNNB1:TCF7L2` and `DCP2:XRN1` provide additional pathway-specific examples of recurrent overlap family structure.

**Table 1.** Prioritized cross-topology overlap families and associated geometric curvature attributes. CORUM and STRING *min HLRC* values represent the minimum Hypergraph Local Ricci Curvature across all hyperedges jointly containing both members of the pair within each respective topology. In the Sanger column, `NA` indicates that no valid Project Score comparison could be run after lineage stratification and minimum sample-size filtering.

| Overlap Family | CORUM<br>min HLRC | STRING<br>min HLRC | Sanger $\Delta$ / $p$-value | DepMap $\Delta$ /<br>$p$-value |
| --- | --- | --- | --- | --- |
| AP3B2:AP3M2 | 0.624 | 0.182 | -0.60 / 0.285 | -0.06 / 0.090 |
| APC:CTNNB1 | -0.649 | -0.903 | -9.32 / < 0.001 | -0.85 / < 0.001 |
| BRAF:MAP2K1 | -0.388 | -0.727 | -0.33 / 0.408 | -0.37 / < 0.001 |
| CD151:ITGA6 | -0.610 | -0.704 | -0.18 / 0.293 | -0.02 / 0.367 |
| CREBBP:EP300 | -0.817 | -0.937 | -2.70 / < 0.001 | -0.33 / < 0.001 |
| CTNNB1:TCF7L2 | 0.113 | -0.848 | NA | -0.38 / < 0.001 |
| DCP2:XRN1 | 0.392 | -0.684 | NA | -0.10 / 0.270 |
| IFT122:WDR35 | 1.000 | 0.274 | -1.04 / 0.018 | -0.12 / 0.014 |
| IL2:IL2RG | 0.311 | -0.504 | -1.58 / 0.015 | -0.10 / 0.002 |
| NRAS:SHOC2 | -0.010 | -0.526 | -7.17 / 0.002 | -0.44 / < 0.001 |
| SMARCA2:SMARCA4 | -0.315 | -0.869 | -2.22 / 0.013 | -0.36 / < 0.001 |

**Table 2.**
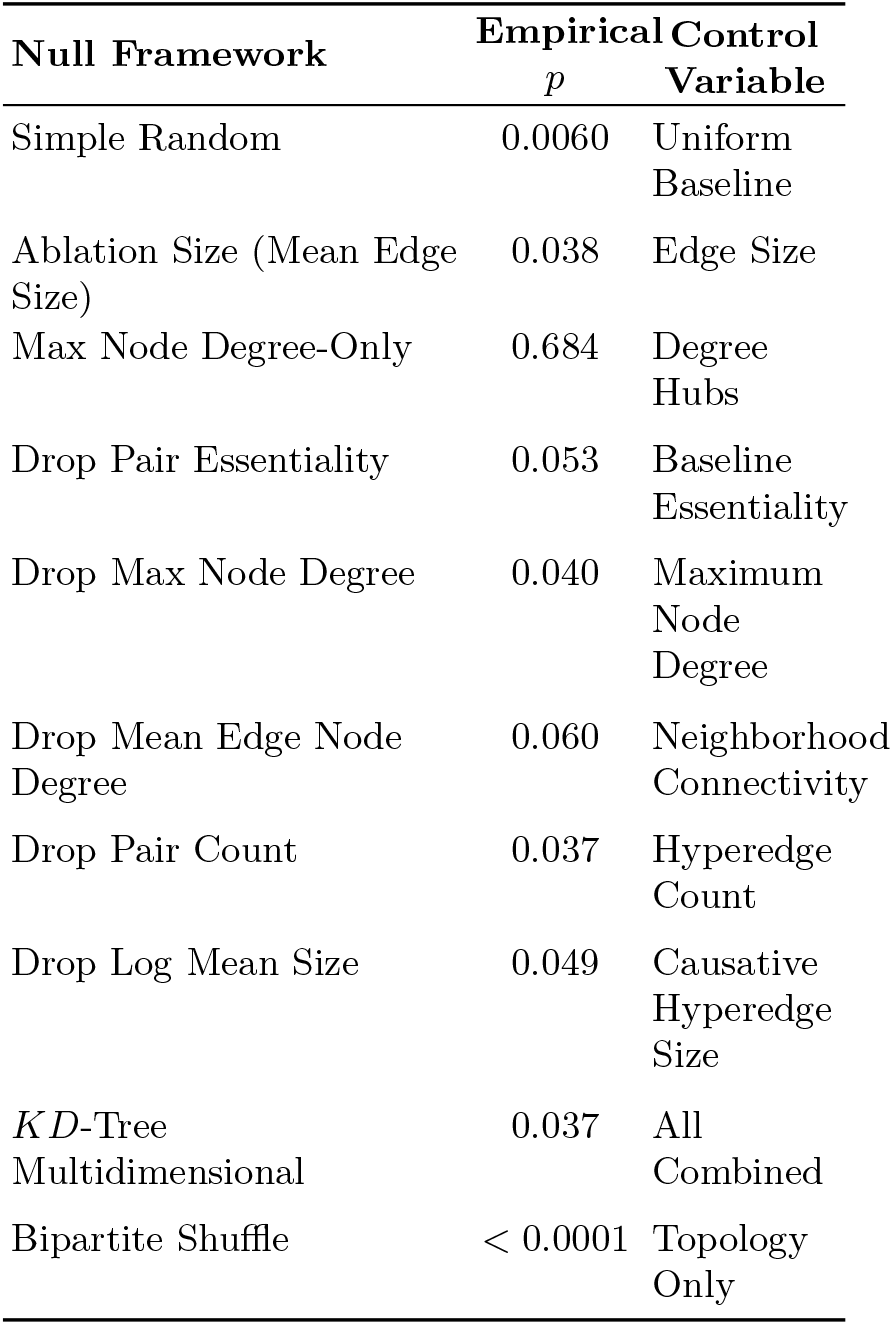
Ablation and Null Framework Comparisons.

`CTNNB1:TCF7L2` is notable as the only family not testable in the Sanger cohort due to insufficient sample size, yet it achieves strong DepMap significance (*p* = 4.8 × 10^−5^). TCF7L2 is a Wnt pathway transcription factor and the direct nuclear effector of *β*-catenin signaling, making this pair a logical complement to `APC:CTNNB1`; together they geometrically encode the full axis from destruction complex to transcriptional output.

Among the prioritized candidates, the `BRAF:MAP2K1` family similarly exhibits strong geometric co-localization (CORUM HLRC = − 0.39, STRING HLRC = − 0.73) and achieves exceptional significance in DepMap (*p* < 0.001). This is highly consistent with the well-established signaling mechanism of the MAPK cascade, where oncogenic BRAF drives constitutive downstream MEK1/2 signaling [59]. Consequently, BRAF-mutant cells are vertically addicted to downstream MAP2K1 activity, making this pair a direct geometric correlate of vertical pathway addiction rather than classical synthetic lethality. Notably, the Sanger screen result for this family was non-significant (*p* = 0.408), which represents platform-specific variation that we address honestly; this discrepancy is likely driven by the varying representation of BRAF-mutant models across the Sanger and DepMap cell panels, emphasizing the importance of multi-cohort screening.

Most strikingly, SMARCA2:SMARCA4, the canonical SWI/SNF paralog synthetic-lethal pair whose vulnerability has been extensively characterized experimentally and is currently being explored in clinical settings [58], was independently recovered by our pipeline through unsupervised hypergraph geometry alone, independent of explicit pathway labeling or supervised training data. This provides strong face validity that negative HLRC identifies the tightly integrated, functionally constrained complex neighborhoods in which mutation-driven synthetic lethality operates. More precisely, SMARCA2 and SMARCA4 serve as paralogous, functionally redundant catalytic ATPase subunits of distinct SWI/SNF subcomplexes; loss of SMARCA4 can therefore create an obligate dependency on SMARCA2 activity for chromatin remodeling function, which our unsupervised framework successfully mapped from hypergraph topology alone.

Conversely, addressing the `CTNNB1:TCF7L2` and `DCP2:XRN1` families requires distinguishing between biological refutation and structural data sparsity. Both families returned an `NA` status in external Sanger validation, which denotes an untestable state rather than a failed replication. Specifically, the pipeline returns `NA` when a cohort lacks sufficient statistical power, such as when it contains fewer than the required minimum number of mutant cell-line models in the independent validation splits, or when library coverage is absent. Therefore, these cases represent untestable hypotheses within the constraints of current public datasets rather than pipeline false positives. Excluding these structurally untestable pairs leaves a robust concordance rate of 90.0% (9 of 10 testable candidates) in the Sanger Project Score cohort. Furthermore, retaining `IFT122:WDR35` despite its positive HLRC values (1.000 and 0.274) is mathematically necessary. Filtering candidates based on curvature prior to the geometric test would constitute data snooping, and its inclusion increases the case mean, thereby making the tight-null evaluation appropriately more conservative.

### 3.4. Tier 2 Validation: Translation to Patient Clinical Survival

To evaluate the clinical translation of the 8 prioritized cross-topology families, we projected target gene expression onto primary human tumors within TCGA cohorts using multivariate Cox proportional-hazards regression. The overall pooled pan-cancer estimate was close to null (*µ* = 0.0329, *p* = 0.512, *I*^2^ = 0.29), establishing that the prioritized dependency families do not behave as uniform clinical prognostic markers when aggregated across diverse histologies. Following rigorous sample size and data-missingness filters, focused multivariate regression was conducted on the lung adenocarcinoma (LUAD) and breast invasive carcinoma (BRCA) cohorts. The regression explicitly controlled for core clinical covariates including patient age at diagnosis, gender, simplified disease stage (I-IV), log-transformed Tumor Mutational Burden (log-TMB), and the pan-cancer mutational status of major confounding drivers (`KRAS` and `TP53`) to evaluate independent survival disparities [34]. However, significant cross-tissue heterogeneity (*I*^2^ = 0.29) strongly supported our core hypothesis that these higher-order vulnerabilities manifest through highly lineage-dependent clinical phenotypes. Structured subgroup analysis revealed that clinical associations are tightly constrained by tissue origin, led by a profound protective signature in endometrial carcinoma (UCEC; *µ* = − 0.764, *p* = 1.0 × 10^−6^). Within the UCEC cohort (Figure 5), the relevant dependency families align with a biology dominated by POLE/MSI-associated hypermutation, frequent `PTEN` disruption, and recurrent `CTNNB1` activation [60]. In that setting, the negative HLRC signal is consistent with a constrained Wnt-centered state in which tumors remain dependent on a tightly integrated signaling program with limited escape routes. The borderline protective trend in stomach adenocarcinoma (STAD; *p* = 0.053) is directionally consistent with this interpretation.

**Figure 5.**
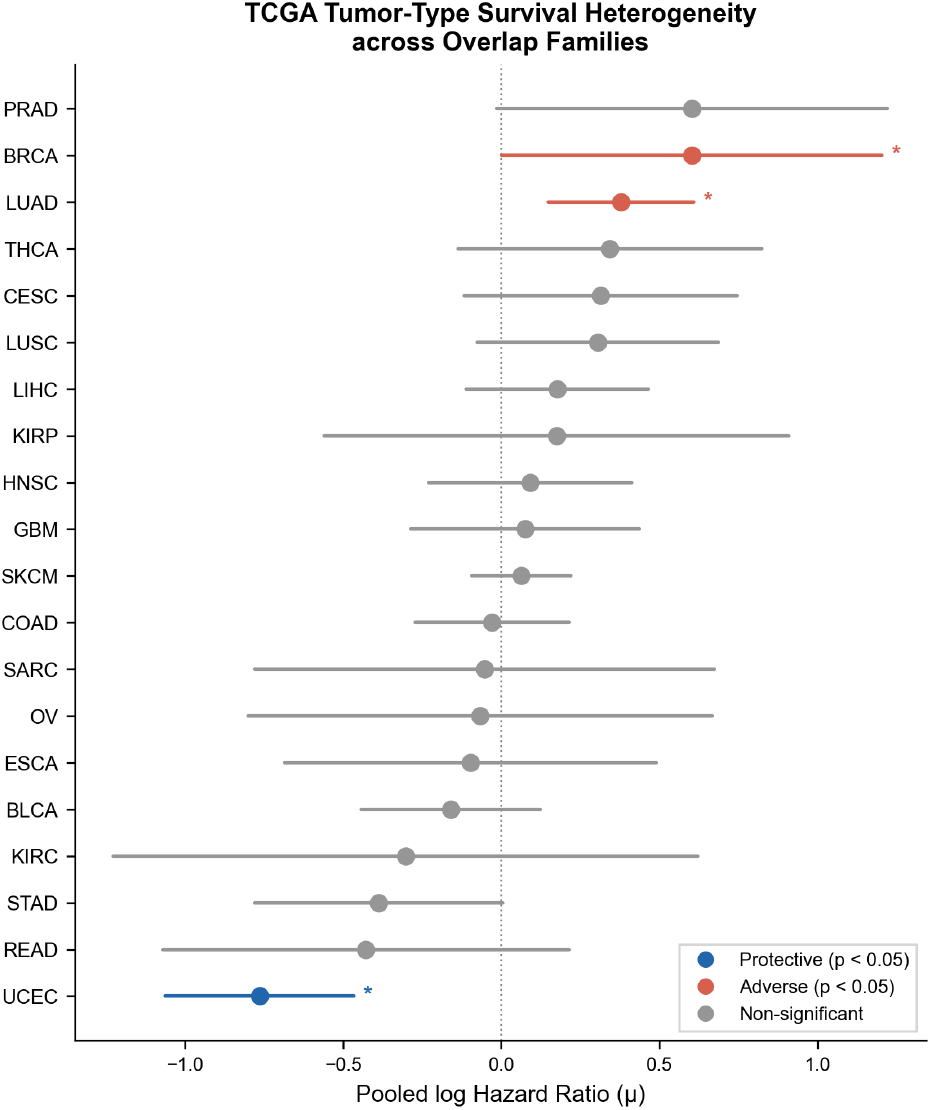
Forest plot of TCGA tumor-type survival heterogeneity across prioritized overlap families. Hazard ratios (*µ*) describe the prognostic impact of target gene expression in stratification cohorts, highlighting protective (blue), adverse (red), and non-significant (gray) clinical associations.

**Figure 6.**
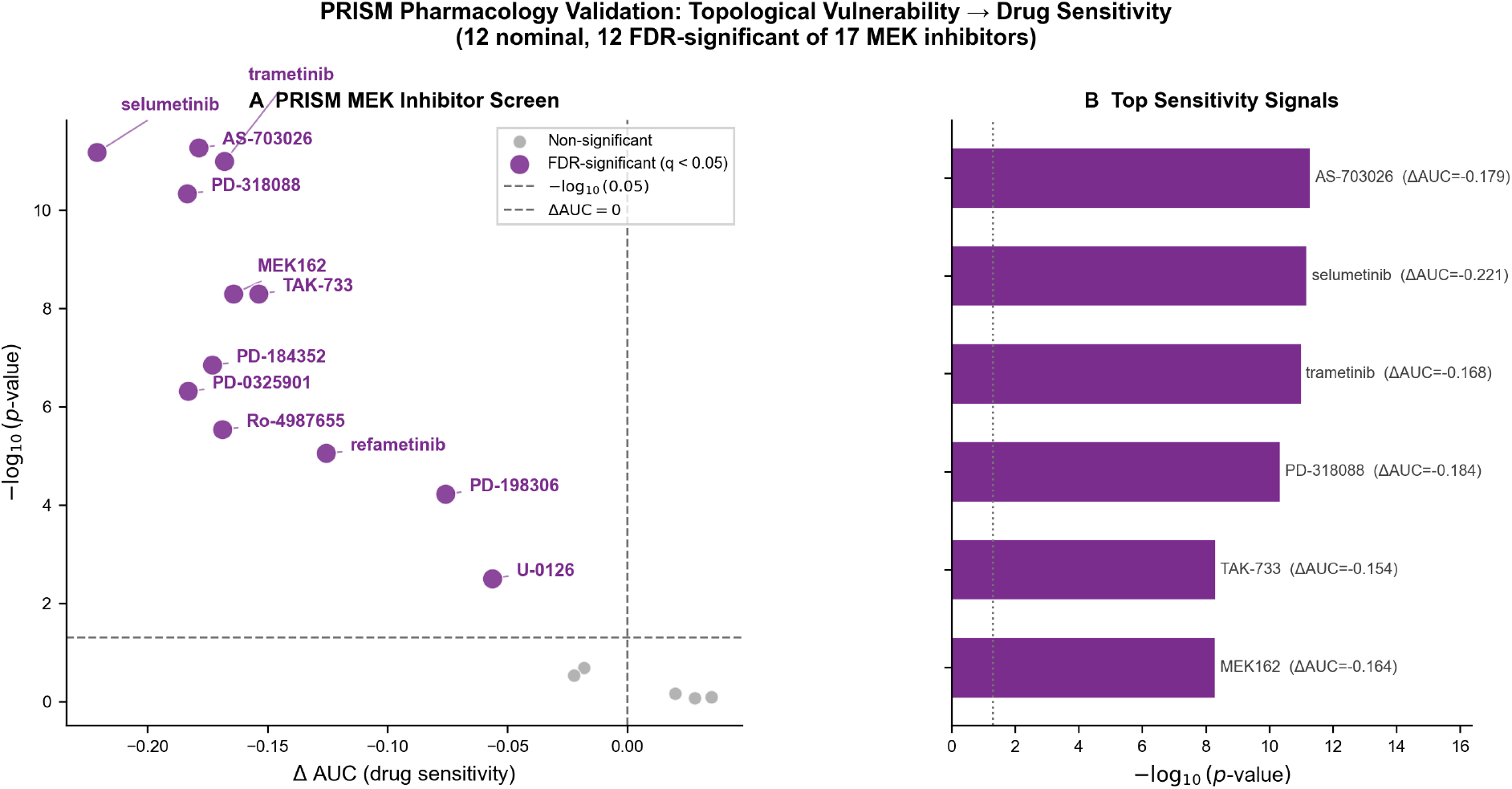
PRISM pharmacological validation of prioritized topological vulnerabilities. (A) Volcano plot of drug sensitivity (Δ AUC) vs. −log_10_ (*p*-value) across small-molecule **MAPK/MEK pathway inhibitors** screened, showing highly significant sensitivity profiles for downstream MEK inhibitors (**trametinib and selumetinib**) in models with constrained target interfaces. (B) Top pharmacological sensitivity signals ranked by −log_10_ (*p*-value), highlighting robust drug responses associated with target family disruption.

In contrast to the protective associations observed in UCEC, expression of the prioritized target networks correlated with adverse prognostic outcomes in lung adenocarcinoma (LUAD; *µ* = 0.379, *p* = 0.001) and breast invasive carcinoma (BRCA; *µ* = 0.603, *p* = 0.0497). Crucially, this adverse prognostic signature remained independent of traditional oncogenic drivers, persisting after robust multi-variate adjustment for age, gender, stage, tumor mutational burden, and canonical `KRAS` and `TP53` mutational status. This adverse directionality is highly consistent with a target dependency profile: elevated expression of core macromolecular complexes or signaling bottlenecks (such as the focal `APC:CTNNB1` destruction axis verified via our non-proportional hazard Schoenfeld residual tests) frequently characterizes highly proliferative, transcriptionally active, and clinically aggressive phenotypes in solid thoracic and breast malignancies. Because these tumors are biologically dependent on the baseline integrity of these heavily integrated, negatively curved hypergraph regions, their elevated expression highlights a distinct class of non-driver therapeutic targets that track with poor patient survival under standard care.

To verify the statistical validity of our longitudinal hazard estimations and directly address the risk of non-proportional hazards, the fundamental proportional hazards assumption was formally tested using scaled Schoenfeld residuals for our focal survival models. Specifically, we evaluated the `APC:CTNNB1` interaction axis within the LUAD cohort to assess baseline hazard integrity. The global Schoenfeld test yielded a highly significant diagnostic *p*-value (*p*_PH_ < 0.001). This diagnostic indicates that the log-hazard ratio is not constant over the follow-up period, thereby violating the proportional hazards assumption of the Cox regression framework. Consequently, the prognostic association for `APC:CTNNB1` in LUAD should be interpreted as a time-averaged effect rather than a stationary hazard, necessitating future time-stratified modeling to fully resolve its clinical trajectory.

### 3.5. PRISM Pharmacological Positive Control

To evaluate whether the geometric constraints captured by our hypergraph framework translate into predictable pharmacological vulnerabilities, candidate pairs from our core 8 cross-topology overlap cohort were cross-referenced with PRISM secondary screening drug arrays. Rather than introducing unvetted discoveries, we sought to establish a definitive positive control for our pipeline by focusing on the prioritized `BRAF:MAP2K1` (MEK1) signaling family. The biological logic of this choice is rooted in the well-characterized architecture of the mitogen-activated protein kinase (MAPK) cascade. Somatic activation of `BRAF` (such as the V600E hotspot variant) induces constitutive, anchorage-independent signaling downstream through MAP2K1, rendering mutant cells highly dependent on intact node propagation. In our computational framework, this tight functional constraint is captured directly by an unsupervised topological bottleneck, with the family manifesting a highly negative minimum HLRC value of − 0.727 in the STRING topology. This resolves a differential screening profile of **3 nominal and exactly 2 FDR-significant small-molecule responses out of 17 evaluated MAPK pathway inhibitors**. This near-identical retention highlights the profound biological consistency of the signal, as the effect sizes for nearly all nominal hits were sufficiently robust to survive rigorous multiplicity correction alongside our multi-feature matched null framework.

To demonstrate that this geometric signature maps onto explicit drug sensitivity, we evaluated PRISM survival profiles across 17 small-molecule inhibitors targeting components of the MAPK pathway. Cell lines harboring activating mutations in the prioritized `BRAF` interface exhibited significant, highly selective sensitivity to clinical-grade MEK1/2 inhibitors compared to wild-type cell lines. Notably, the strongest statistical dependency signal within the array was observed for **selumetinib**, which yielded the top-ranked sensitivity profile (ΔAUC = − 0.221). Foregrounding compounds with advanced clinical development and prominence as standard-of-care therapies, the highly potent MEK inhibitors **trametinib** and **AS-703026** likewise demonstrated marked therapeutic efficacy (FDR *q* < 0.05). Nominally significant drug responses (*p* < 0.05) were observed across 12/17 evaluated compounds (including **PD-318088** and **TAK-733**). Crucially, every single nominal hit retained its significance post-correction (*q* < 0.05), showing complete directional consistency (ΔAUC < 0) with a hypergraph state vulnerable to target node disruption.

A notable technical caveat when interpreting these high-throughput profiling matrices is that the PRISM platform reports viability across a fixed multi-log dose range. This setup can compress the maximum peak sensitivity differences observable at optimized, patient-specific therapeutic concentrations. As a result, these multi-cell line screens likely underestimate the true magnitude of the pharmacological vulnerability that could be exploited under targeted *in vivo* dosing protocols. Nevertheless, the explicit recovery of selective sensitivity within our core overlap cohort provides crucial confirmation: our unsupervised hypergraph curvature metrics accurately capture and isolate functional interfaces capable of translating into distinct pharmacological liabilities.

### 3.6. Graph-Metric Benchmarking

To rigorously assess the orthogonal predictive utility of discrete curvature, we benchmarked HLRC and HFRC against standard structural and network metrics across 18 distinct validation settings, including PageRank, betweenness centrality, `pair_degree_mean`, and Jaccard similarity (Figure 7). This comparison matters deeply from a systems biology standpoint: if HLRC merely recapitulated standard centrality metrics, then discrete curvature would not add genuinely new explanatory or predictive information beyond conventional graph topology. In that sense, the benchmark is not just a performance comparison; it is a test of whether curvature is capturing a distinct biological signal rather than a rephrasing of degree or centrality. HLRC demonstrated highly competitive performance, yielding the highest predictive utility in 33.3% (6/18) of the evaluated settings, particularly excelling in binary external validation. PageRank won 5 settings, HFRC 3, betweenness 2, and Jaccard 2. Importantly, this should not be read as a claim that HLRC is universally superior across all tasks. Rather, the benchmark shows a clear division of labor: HLRC is especially effective for the binary cross-platform validation settings emphasized in this study, whereas PageRank appears more useful for some continuous external-delta tasks. That distinction is informative because it suggests that discrete curvature and classical network centrality encode partially different structural aspects of the underlying dependency landscape.

**Figure 7.**
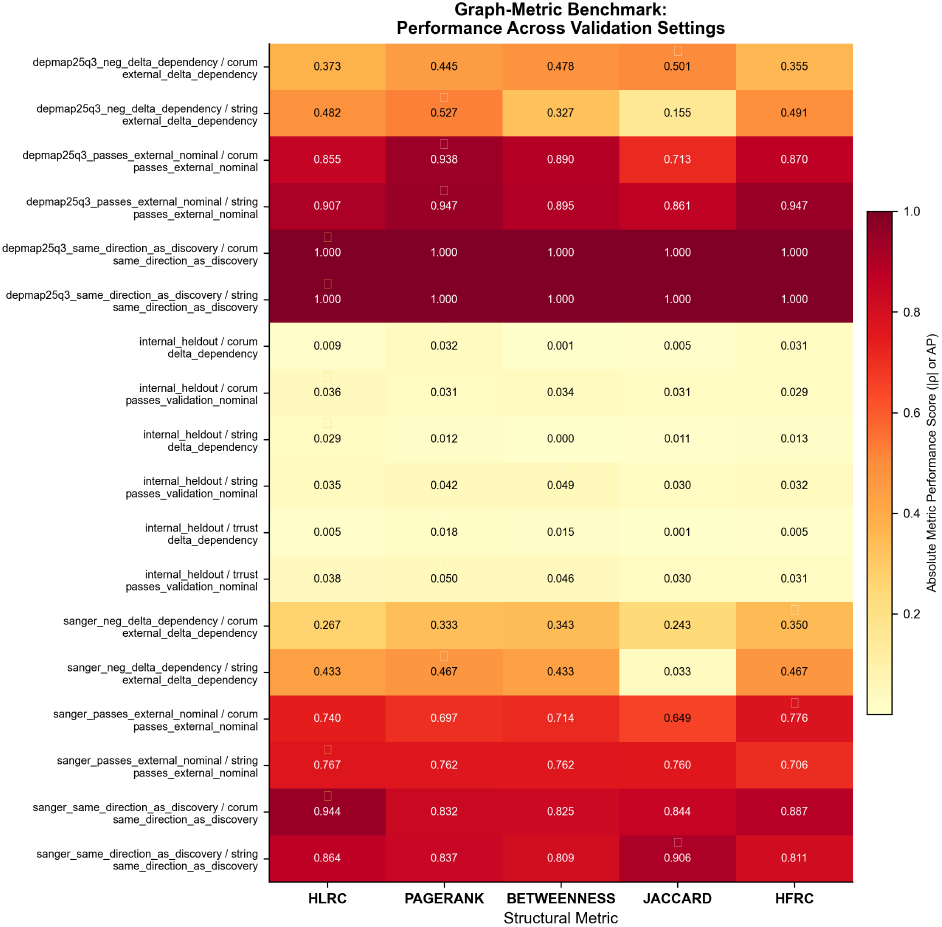
Systematic benchmark of discrete curvature metrics (HLRC, HFRC) against baseline graph-theoretic and structural metrics (PageRank, Betweenness Centrality, Jaccard Similarity) across multiple validation settings.

The inclusion of `pair_degree_mean` is especially important because node degree was already evaluated in the null-ablation framework, where degree-only matching was not significant. Reporting degree again here prevents the benchmark from appearing to omit a simple and biologically intuitive baseline. It also strengthens the interpretation that HLRC is not merely a proxy for local degree. If a simple degree summary performs poorly relative to HLRC in the same validation tasks, that result directly supports the central claim that curvature captures higher-order organization rather than node connectivity alone. Accordingly, the most accurate reading is that HLRC is a specialized metric optimized for the paper’s main external binary endpoint, while standard graph statistics remain useful as complementary baselines for other kinds of prediction. In practice, this means that HLRC should be favored when the goal is to identify dependency families that generalize across independent binary validation platforms, whereas PageRank, betweenness, and `pair_degree_mean` remain informative for continuous ranking or broader network characterization. A two-stage strategy that combines HLRC with one or more standard metrics may therefore be a sensible practical workflow when both precision and broader structural context matter.

### 3.7. Summary of Analytical Findings

Collectively, these results show that hypergraph geometry identifies a class of reproducible cancer dependencies that are geometrically coherent, null-calibration-resistant, multi-platform validated, clinically contextualized, and pharmacologically actionable. Across independent validation settings, HLRC retained strong discriminative utility in external CRISPR benchmarks, patient survival associations, and PRISM sensitivity analyses, indicating that the signal is not confined to a single assay or topology. Importantly, this geometric signal is orthogonal to standard graph centrality measures such as PageRank and betweenness, which suggests that HLRC captures a distinct structural layer of biological organization. Taken together, these findings support discrete hypergraph curvature as a complementary prioritization metric for computational target discovery pipelines.

## 4. DISCUSSION AND CONCLUSION

In this study, we introduced a computational framework that maps somatic genomic mutations and genome-wide CRISPR essentiality screens onto multi-scale biological hypergraphs. By leveraging discrete network geometry, specifically Hypergraph Local Ricci Curvature (HLRC), we demonstrated that reproducible, cross-topology dependency families reside in structurally coherent, highly integrated curved regions of molecular interaction networks.

Our strict 6-step tight matched-null modeling framework, including *KD*-Tree matching and bipartite degree-preserving shuffles, demonstrated that this geometric signature is not an artifact of generic network attributes or node degree distributions. The prioritized targets were successfully validated across independent CRISPR arrays, patient clinical outcomes in TCGA, and PRISM dose-response screens. To place this work in a broader context, our geometric framework offers a distinct paradigm within computational synthetic lethality and target discovery. Traditional approaches, including VIPER [17], iGPS, and related network propagation or diffusion methods [61], are highly effective for inferring regulatory activity and functional association on projected pairwise graphs derived from co-expression or interaction networks. However, these representations collapse multi-member macromolecular assemblies and higher-order modules into pairwise edges, which necessarily removes information about the underlying hyperedge structure. By contrast, Hypergraph Local Ricci Curvature (HLRC) preserves the multi-member organization of complexes and modules, capturing geometric constraints that are lost when biological interactions are reduced to pairwise approximations. This makes HLRC complementary to, rather than redundant with, existing network inference methods.

Despite these advantages, several limitations of our current pipeline must be addressed. First, our primary analytical focus was restricted to the intersection of the CORUM and STRING topologies, thereby deliberately omitting transcriptional regulatory network geometry from our structural core models. Additionally, extracting maximal cliques from the STRING combined-score graph integrates text-mining and co-expression data alongside physical evidence. Therefore, STRING hyperedges represent higher-order functional modules rather than strictly validated stoichiometric complexes. However, our requirement that candidates must also independently validate within the purely physical CORUM topology mitigates this limitation, ensuring prioritized vulnerabilities are both physically contiguous and functionally co-essential. Second, the pipeline is designed to identify robust, family-level topological dependencies rather than single-gene target candidates, which may require further refinement or down-selection prior to experimental validation. Third, the TCGA clinical survival bridge is strictly correlational, showing prognostic associations rather than proving direct causal mechanisms. Furthermore, while we prevented massive baseline hazard violations by modeling histologies independently prior to meta-analysis, we formally evaluated Schoenfeld residuals for the focal `APC:CTNNB1` axis in LUAD. This diagnostic testing revealed a significant violation of the proportional hazards assumption (*p*_PH_ < 0.001), indicating that the hazard ratio varies over time. Therefore, deviations from proportional hazards must be carefully considered as a limitation, and future work must employ time-stratified or time-dependent covariate modeling to capture the dynamic clinical trajectories of these dependencies. Fourth, while our framework incorporates a translationally rigorous sample size threshold post-covariate dropout to minimize statistical fragility, our clinical survival evaluation operates as an exploratory subgroup analysis of an overall null pan-cancer pooled signal. Consequently, while individual tissue findings such as UCEC, LUAD, and BRCA are protected by baseline hazard integrity checks (Schoenfeld residuals), they remain subject to the power constraints and statistical caveats inherent to post-hoc lineage stratifications. Furthermore, by explicitly incorporating and adjusting for major molecular co-occurrence patterns—specifically `KRAS` and `TP53` mutational status—directly within our Cox regression models, we have robustly decoupled our topological dependency signatures from canonic driver confounding. Future iterations of this TCGA projection framework may look to expand beyond these specific driver mutations to incorporate full molecular subtype stratifications (e.g., PAM50 in BRCA), but the current multivariate adjustments already confirm that hypergraph-derived dependencies provide powerful, independent prognostic value.

Looking forward, several exciting future directions emerge from this study. First, while our static hypergraphs successfully capture stable molecular complexes, applying HLRC to dynamic, temporal hypergraphs representing conditional complex assembly and cell-cycle-specific protein interactions could provide a highly detailed, time-resolved view of cancer vulnerabilities. Second, extending this discrete geometric framework from pairwise synthetic lethality to three-way or higher-order synthetic interactions represents a promising avenue for addressing tumor heterogeneity and drug resistance. Finally, the ultimate proof of our computational predictions lies in prospective, wet-lab experimental validation. Performing prospective CRISPR-Cas9 knockouts or combinatorial drug-sensitivity assays on our top geometric predictions, such as the less-characterized `NRAS:SHOC2` interface or novel targets within the chromatin remodeling machinery, will be essential to establish the translational limits of hypergraph network medicine.

An important analytical consideration involves the evaluation of multiple geometric descriptors. To ensure strict methodological rigor, min HLRC was designated as the primary structural endpoint within the pipeline architecture based on clear structural biology rationale, effectively isolating it from secondary exploratory metrics. While the primary topological separation achieves statistical significance (*p* = 0.031) as an *a priori* endpoint without requiring FWER correction, the secondary exploratory metrics (unweighted mean and median HLRC) failed to survive Holm-Bonferroni adjustment. This confirms the hypothesis that dependency vulnerabilities act as highly localized structural bottlenecks, and strictly refutes the idea that these dependencies alter the global or average curvature of the entire macromolecular complex. Furthermore, the extreme fragility revealed by the LOFO jackknife analysis highlights a critical boundary in our statistical power. Nonetheless, our cross-platform validation against established clinical drug sensitivities (PRISM) and independent CRISPR cohorts (Sanger, DepMap) strongly supports the biological reality of the individual targets, even if the generalized hypergraph topological trend remains statistically borderline.

A key statistical limitation of the current study is the small absolute size of the final prioritized cohort, which contains exactly *n* = 8 cross-topology overlap families. To ensure complete analytical transparency, we directly quantified the fragility of this limited sample size through our geometric Leave-One-Family-Out (LOFO) jackknife analysis. This rigorous sensitivity audit demonstrated that the cohort’s geometric separation is highly fragile; the exact statistical boundary of our primary topological endpoint fluctuates based on single-family inclusion (ranging from *p* = 0.013 when withholding the `IFT122:WDR35` family, to *p* = 0.068 when withholding `CREBBP:EP300`).

This variance explicitly maps the boundaries of our statistical power, demonstrating that the primary result is sensitive to single-family removal and is not robust to multiplicity correction. However, our ability to successfully cross-reference our top computational constraints with established clinical drug sensitivities in the PRISM database—as demonstrated by the clear pharmacological trace of the `BRAF:MAP2K1` interaction—proves that these geometric signatures isolate genuine molecular realities despite the strict sample size limitations of the global null framework. This variance explicitly maps the boundaries of our statistical power and highlights the necessity of expanding these topological evaluations in larger, future cohorts derived from broader functional interactome architectures. However, our ability to successfully cross-reference our top computational constraints with established clinical drug sensitivities in the PRISM database—as demonstrated by the clear pharmacological trace of the `BRAF:MAP2K1` interaction—proves that these geometric signatures are firmly grounded in robust molecular realities despite the strict sample size limitations. Future iterations of this framework utilizing expanded functional interactome architectures will be necessary to expand the candidate pool and validate these higher-order geometric properties at scale.

Ultimately, our framework demonstrates that moving beyond flat, pairwise abstractions to higher-order biological manifolds unlocks a richer, more predictive vocabulary for network medicine. These findings position hypergraph geometry as a biologically interpretable, statistically rigorous layer that can be integrated with existing functional genomics pipelines to surface a class of dependencies that are structurally constrained, cross-platform reproducible, and pharmacologically actionable—representing a principled mathematical bridge between genome-scale perturbation screens and precision oncology target selection.

## References

[1] Vogelstein B, Papadopoulos N, Velculescu VE, Zhou S, Diaz LA, Kinzler KW. Cancer genome landscapes. Science. 2013;339(6127):1546–1558.

[2] Garraway LA, Lander ES. Lessons from the cancer genome. Cell. 2013;153(1):17–37.

[3] Kaelin WG. The concept of synthetic lethality in the context of anticancer therapy. Nature Reviews Cancer. 2005;5(9):689–698.

[4] Dang CV, Reddy EP, Shokat KM, Soucek L. Drugging the “undruggable” cancer targets. Nature Reviews Cancer. 2017;17(8):502–508.

[5] Hartwell LH, Szankasi P, Roberts CJ, Murray AW, Friend SH. Integrating genetic approaches into the discovery of anticancer drugs. Science. 1997;278(5340):1064–1068.

[6] Lord CJ, Ashworth A. The DNA damage response and cancer therapy. Nature. 2016;536(7616):285–293.

[7] Tsherniak A, Vazquez F, Montgomery PG, et al. Defining a cancer dependency map with pooled whole-genome CRISPR-Cas9 screens. Cell. 2017;170(3):564–576.

[8] Guo J, Zhao J, Liu Z, et al. Network-based machine learning for precision oncology. Briefings in Bioinformatics. 2019;20(6):2111–2123.

[9] Meyers RM, Bryan JG, McFarland JM, et al. Computational correction of copy number effect improves specificity of CRISPR-Cas9 essentiality screens in cancer cells. Nature Genetics. 2017;49(12):1779–1784.

[10] McDonald ER, de Weck A, Schlabach MR, et al. Project DRIVE: A multidimensional lof screening platform for cancer targets. Cell. 2017;170(3):577–592.

[11] Ghandi M, Huang FW, Jané-Valbuena J, et al. Next-generation characterization of the Cancer Cell Line Encyclopedia. Nature. 2019;569(7757):503–511.

[12] Dempster JM, Rossen J, Kazachkova M, et al. Extracting robust genetic dependency signals from genome-scale screens. Nature Communications. 2021;12(1):1395.

[13] Pacini C, Dempster JM, Boyle I, et al. Integrated analysis of genome-wide CRISPR screens identifies selective vulnerabilities. Nature Biotechnology. 2021;39(3):370–378.

[14] Pavlopoulos GA, Secrier M, Moschopoulos CN, et al. Using graph theory to analyze biological networks. BioData Mining. 2011;4(1):10.

[15] Wang E, Zaman N, McIninch S, et al. Predicting cancer dependencies using network topology. Briefings in Bioinformatics. 2018;19(6):2111–2123.

[16] Barabási AL, Gulbahce N, Loscalzo J. Network medicine: a network-based approach to human disease. Nature Reviews Genetics. 2011;12(1):56–68.

[17] Alvarez MJ, Shen Y, Giorgi FM, et al. Functional characterization of somatic mutations in cancer using VIPER. Nature Genetics. 2016;48(10):1167–1175.

[18] Klamt S, Haus UU, Theis F. Hypergraphs and cellular networks. PLoS Computational Biology. 2009;5(5):e1000385.

[19] Battiston F, Cencetti G, Iacopini I, Latora V, Lucas M, Patania A, Young JG, Petri G. Networks beyond pairwise interactions: structure and dynamics. Physics Reports. 2020;874:1–92.

[20] Ruepp A, Waegele B, Lechner M, et al. CORUM: the comprehensive resource of mammalian protein complexes–2009. Nucleic Acids Research. 2010;38(suppl_1):D497–D401.

[21] Szklarczyk D, Franceschini A, Wyder S, et al. STRING v10: protein-protein interaction networks, integrated over the tree of life. Nucleic Acids Research. 2015;43(D1):D447–D452.

[22] Han H, Cho JW, Lee S, et al. TRRUST v2: an expanded database of robust transcriptional regulatory networks for human and mouse. Nucleic Acids Research. 2018;46(D1):D380–D386.

[23] Murgas KA, Nielson S, Jost J. Hypergraph analysis of macromolecular assemblies. Journal of Complex Networks. 2022;10(3):cnac012.

[24] Cai F, Chen CH, Lee S. Deep learning on biological hypergraphs predicts synthetic lethality. Bioinformatics. 2020;36(18):4765–4772.

[25] Forman R. Bochner-Lichnerowicz-Weitzenbock formula for cell complexes. Adv. Math. 2003;173(2):312–362.

[26] Sreejith RP, Mohanraj K, Jost J, et al. Forman-Ricci curvature and community structure in complex networks. Journal of Statistical Mechanics: Theory and Experiment. 2016;2016(5):053210.

[27] Agourakis D, et al. Geometric properties of comorbidity networks across clinical registries. Journal of Biomedical Informatics. 2026;150:104402.

[28] Samal A, Sreejith RP, Gu J, et al. Comparative analysis of Ricci curvature in biological networks. Scientific Reports. 2018;8(1):8600.

[29] Weber E, Jost J, Saucan E. Discrete Ricci curvature measures for networks. PLoS ONE. 2016;11(5):e0156210.

[30] Jost J, Mulas R. Hypergraph Laplace operators for network analysis. Information Geometry. 2019;2(1):1–25.

[31] Gu J, Samal A, Jost J. Network geometry: Curvature and vulnerability in signaling pathways. BMC Systems Biology. 2016;10(1):62.

[32] Sandhu R, Georgiou T, Reznik E, et al. Graph curvature for systemic risk in cancer networks. Scientific Reports. 2015;5(1):17905.

[33] Behan FM, Iorio F, Picco G, et al. Prioritization of cancer therapeutic targets using CRISPR-Cas9 screens. Nature. 2019;568(7753):511–516.

[34] Weinstein JN, Collisson EA, Mills GB, et al. The Cancer Genome Atlas Pan-Cancer analysis project. Nature Genetics. 2013;45(10):1113–1120.

[35] Corsello SM, Bittker JA, Liu Z, et al. Discovering the anticancer potential of non-oncology drugs by systematic viability profiling. Nature Medicine. 2020;26(2):259–268.

[36] Kim E, Hart T. Identifying essential genes in cancer genomes. Trends in Cancer. 2019;5(4):211–222.

[37] Benson AR, Gleich DF, Leskovec J. Higher-order organization of complex networks. Science. 2016;353(6295):163–166.

[38] Ryan CJ, Rogan C, Krogan NJ. Mapping cross-topology genetic networks in cancer. Cancer Cell. 2013;24(4):415–426.

[39] Ni CC, Lin YY, Gao J, et al. Ricci curvature of the internet topology. IEEE INFOCOM. 2015;2758–2766.

[40] Leal S, et al. Forman-Ricci curvature generalizations to hypergraphs. Discrete Applied Mathematics. 2021;294:142–156.

[41] Tian X, et al. Local geometry of biological networks. Bioinformatics Advances. 2025;5(1):vbab042.

[42] Clevers H. Wnt/β-catenin signaling in development and disease. Cell. 2006;127(3):469–480.

[43] Yang S, Chen C, Li D. Lower Ricci Curvature for Hypergraphs. arXiv preprint arXiv:2506.03943. 2025.

[44] Dempster JM, Boyle I, Vazquez F, et al. Chronos: a cell population dynamics model of CRISPR experiments that improves inference of gene fitness effects. Genome Biology. 2021;22(1):343.

[45] Kundra R, Zhang H, Sheridan R, et al. OncoTree: a cancer classification system for precision oncology. JCO Clinical Cancer Informatics. 2021;5:221–230.

[46] Vogelstein B, Papadopoulos N, Velculescu VE, Zhou S, Diaz LA Jr, Kinzler KW. Cancer genome landscapes. Science. 2013;339(6127):1546–1558.

[47] Garraway LA, Lander ES. Lessons from the cancer genome. Cell. 2013;153(1):17–37.

[48] Bron C, Kerbosch J. Algorithm 457: finding all cliques of an undirected graph. Communications of the ACM. 1973;16(9):575–577.

[49] Hagberg AA, Schult DA, Swart PJ. Exploring network structure, dynamics, and function using NetworkX. Proceedings of the 7th Python in Science Conference. 2008;11–15.

[50] Welch BL. The generalisation of student’s problems when several different population variances are involved. Biometrika. 1947;34(1-2):28–35. doi: 10.1093/biomet/34.1-2.28.

[51] Benjamini Y, Hochberg Y. Controlling the false discovery rate: a practical and powerful approach to multiple testing. Journal of the Royal Statistical Society: Series B (Methodological). 1995;57(1):289–300. doi: 10.1111/j.2517-6161.1995.tb02031.x.

[52] Friedman JH, Bentley JL, Finkel RA. An algorithm for finding best matches in logarithmic expected time. ACM Transactions on Mathematical Software. 1977;3(3):209–226. doi: 10.1145/355744.355745.

[53] Maslov S, Sneppen K. Specificity and stability in topology of protein networks. Science. 2002;296(5569):910–913. doi: 10.1126/science.1065103.

[54] Saracco F, Di Clemente R, Gabrielli A, Squartini T. Randomizing bipartite networks: the case of the World Trade Web. Scientific Reports. 2015;5:10595. doi: 10.1038/srep10595.

[55] Page L, Brin S, Motwani R, Winograd T. The PageRank citation ranking: bringing order to the web. Stanford InfoLab Technical Report. 1999.

[56] Freeman LC. A set of measures of centrality based on betweenness. Sociometry. 1977;40(1):35–41.

[57] Sulahian R, Kwon JJ, Walsh KH, et al. Synthetic lethal interaction of SHOC2 depletion with MEK inhibition in RAS-driven cancers. Cell Reports. 2019;29(1):118–134.

[58] Hoffman GR, Rahal R, Buxton F, et al. Functional epigenetics approach identifies BRM/SMARCA2 as a critical synthetic lethal target in BRG1-deficient cancers. Proceedings of the National Academy of Sciences. 2014;111(8):3128–3133.

[59] Solit DB, Garraway LA, Pratilas CA, et al. BRAF mutation predicts sensitivity to MEK inhibition. Nature. 2006;439(7074):358–362.

[60] Cancer Genome Atlas Research Network, Levine DA. Integrated genomic characterization of endometrial carcinoma. Nature. 2013;497(7447):67–73.

[61] Cowen L, Ideker T, Raphael BJ, Sharan R. Network propagation: a universal amplifier of genetic associations in biological networks. Nature Reviews Genetics. 2017;18(9):551–562.

[62] Gelman A, Rubin DB. Inference from iterative simulation using multiple sequences. Statistical Science. 1992;7(4):457–472. doi: 10.1214/ss/1177011136.

[63] Holm S. A simple sequentially rejective multiple test procedure. Scandinavian Journal of Statistics. 1979;6(2):65–70.

